# Human endogenous retrovirus activation contributes to biliary atresia pathogenesis through re-education of resident macrophages

**DOI:** 10.1101/2022.03.11.483921

**Authors:** Jianpeng Sheng, Junlei Zhang, Yaxing Zhao, Jinyuan Song, Jianghui Tang, Xun Wang, Yongtao Ji, Jiangchao Wu, Taohong Li, Hui Zhang, Vincent Tano, Sarah Raye Langley, Xueli Bai, Tingbo Liang

## Abstract

Biliary atresia (BA) is a life-threatening neonatal fibro-inflammatory disease characterized by hepatic fibrosis, cirrhosis, and end-stage liver failure. BA is also the most frequent indication of pediatric liver transplantation globally. Despite the devastating condition of BA, the pathogenesis mechanism is unknown. Viral infection has been suggested to be associated with BA, but definitive evidence to support this hypothesis is not available. To elucidate the virus-associated pathogenesis mechanism of BA and to understand the immune ecosystem, we performed single-cell transcriptomic and proteomic profiling of BA livers. We detected human endogenous virus (HERV) in infants with BA and their parents. HERV was mainly found in FOLR2^+^ resident macrophages, T cells, and NK cells. In addition, HERV activation re-educated the fetal-derived FOLR2^+^ resident macrophages, and reactive oxygen species scavenging neutrophil recruitment was impaired in patients with BA and HERV^+^, due to FOLR2^+^ resident macrophage re-education. Furthermore, we showed depletion of FOLR2^+^ resident macrophage and N-acetylcysteine treatment could rescue the liver damage in BA. Overall, our study revealed the HERV-associated immunopathology mechanism of BA. These results contribute to potential diagnosis and immunotherapy strategies for BA.

## Introduction

Biliary atresia (BA) is a devastating neonatal fibro-inflammatory disease characterized by the obstruction of the extrahepatic biliary tract and the absence of normally branching intrahepatic ducts ^1^. Hepatic fibrosis, cirrhosis, and end-stage liver failure are the common fatal consequences of progressive cholestatic jaundice if untreated timely. Kasai portoenterostomy is the classic operation for the condition of congenital biliary atresia performed on infants to allow for bile drainage ^2^. BA is also the most frequent indication of pediatric liver transplantation globally. The development of curative treatment, to alleviate the need for surgical intervention, has been hindered due to its complexity and lack of understanding of the disease etiology and pathogenesis.

The hypothesis that a virus plays a pivotal role in BA pathogenesis, put forward by Landing in 1974 ^3^, sheds light on the emergence of virus candidates for the initiation of BA, such as cytomegalovirus (CMV), Epstein-Barr virus (EBV), rotavirus, reovirus, etc. CMV is by far the best-investigated virus in terms of its relationship to human BA, unfortunately, the reported results have been inconclusive ^4–6^. Although some cases have demonstrated there is a connection between EBV and BA, weak evidence was collected from previous studies to support the correlation ^7–9^. Rates of rotavirus prevalence in BA were noted to fluctuate for both rotavirus types A and C based on the polymerase chain reaction (PCR) experiment ^10–12^. Likewise, the hypothesis of reovirus responsible for the development of BA was overturned by a follow-up study ^3, 13^. In line with the mentioned studies, there is also no evidence supporting viral infections triggering BA in a systematic review with 19 studies on 16 kinds of viruses ^14^. Although many of these findings have been established, there are potential pitfalls of the applied method, PCR, which is highly sensitive and requires a considerable number of samples along with a strictly adjusted and finely tuned system to reduce false positives and obtain reliable and consistent results ^15^. Taken together, the viral infection underlying BA pathogenesis is likely, but the relationship has not been determined.

Recent advances in high-dimensional single-cell modalities provide exciting new opportunities to understand cellular and molecular diversity in healthy tissues and disease. Single-cell RNA sequencing (scRNA-seq) has the potential to enable gene expression profiling at the level of the individual cell ^16^. In parallel, cytometry by time of flight (CyTOF) enable simultaneous single-cell measurement of more than 50 surface and intracellular proteins by combining metal isotope-labeled antibodies with mass spectrometry detection ^17^. In a pilot study, Wang et al characterized a comprehensive liver immune landscape to understand the pathogenesis and therapeutics for BA by single-cell RNA profiling ^18^, however, single-cell proteomic profiling of BA liver hasn’t been performed yet and the virus-associated pathogenesis mechanism is not addressed. Therefore, we carried out a multi-omics study to decipher the contribution of pathological immune response caused by viral components at the single-cell resolution.

To map the immune response of the BA ecosystem at the transcriptional and proteomic levels, we performed single-cell RNA sequencing (scRNA-seq), and cytometry by time of flight (CyTOF) on the liver biopsies from a cohort of BA patients. We discovered human endogenous virus activation was detected in the BA patients and their parents ^19^ and validated through immunohistochemistry (IHC), multiplex IHC (mIHC), and ELISA. Human endogenous retrovirus (HERV) was mainly found in FOLR2^+^ resident macrophages, T cells, and NK cells. In addition, we found the virus infection re-educated the fetal-derived FOLR2 resident macrophage. Although neutrophil was not the direct target of HERV, we found^+^ SOD1/2^+^ neutrophils mediated ROS scavenging was impaired in BA due to FOLR2^+^ resident macrophage re-education, and depletion of FOLR2^+^ resident macrophage could rescue the liver tissue damage in BA. To summarize, our results reveal novel mechanistic insights into the pathogenesis of viral-associated immunological perturbations in BA and shed light on the potential diagnosis and immunotherapy strategies for BA.

## Results

### A Single-Cell Transcriptomic and Proteomic Atlas of Biliary Atresia Ecosystems

To generate a comprehensive immune cell atlas reflecting cellular and systemic adaptations resulting from virus infection, we integrated scRNA-seq and CyTOF tensor analysis of liver biopsies collected from a 12-patient cohort (Fig 1A and B). All the patients enrolled had not gone through Kasai portoenterostomy and were within the ages of 3.9 to 9.4 months. In addition, a similar number of male (5) and female patients (7) were selected (Fig. 1A). Furthermore, important disease indicators were also listed (Fig. 1A), such as alanine aminotransferase (ALT) and aspartate aminotransferase (AST). The detailed clinical information was listed in Table S1.

**Figure 1.**
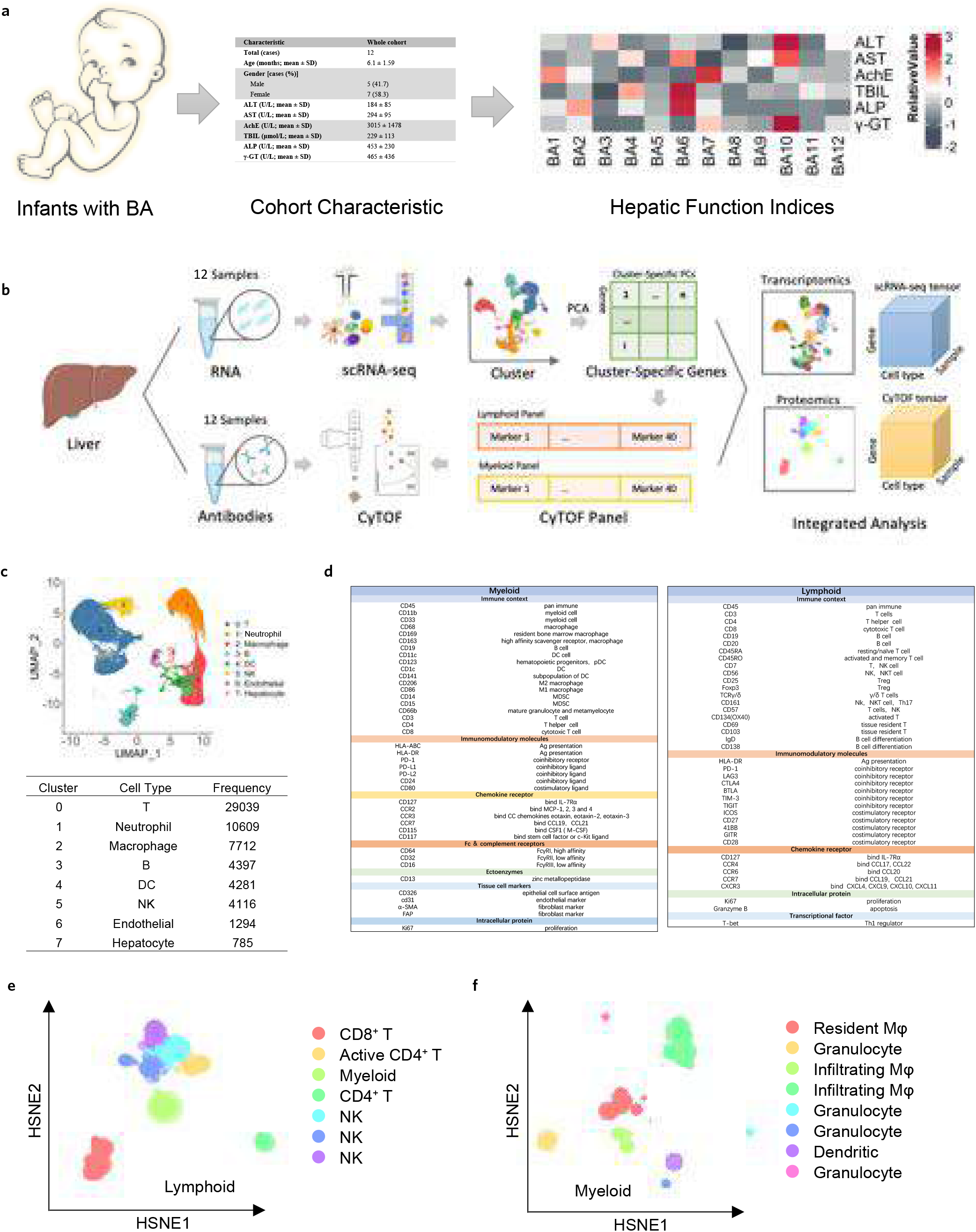
Single-cell transcriptomic and proteomic profiling of the BA ecosystem. A. BA patients’ enrolment scheme. 12 BA patients prior to Kasai portoenterostomy were selected. Patients’ age, sex, and major clinical parameters were shown. B. BA sample analysis pipeline. 12 patients’ samples were processed into single cells for scRNA-seq, followed by cell cluster annotation. Typical markers for each cell cluster were shared for the lymphoid and myeloid panel used for Cy-TOF analysis. 12 patients’ samples were processed into single cells for Cy-TOF analysis and cell cluster definition. Both scRNA-seq and Cy-TOF data went through tensor analysis. C. 8 major cell populations were defined by scRNA-seq analysis, shown by UMAP. And cell count for each population was also calculated after quality control. D. Markers included in the lymphoid and myeloid panel were listed based on their functional category. Many markers were shared with feature genes for each cell clustered defined by scRNA-seq analysis and the remaining markers were classical immune markers. E and F. Major lymphoid and myeloid cell populations defined by HSNE analysis based on CyTOF results were shown. Each colour was annotated to one major cell population.

We ultimately acquired 31,708 high-quality cells (minimum of 200 genes, maximum of 6000 genes, and < 10% mitochondrial reads per cell) from 12 samples using the 10X genomics scRNA-seq platform. The high-quality cells were integrated to eliminate batch effects using the Seurat package ^20–22^. To explore the cellular composition, we distinguished cell types by graph-based clustering based on the RNA expression of cluster-specific markers and visualized by Uniform Manifold Approximation and Projection (UMAP) ^23^, a novel manifold learning technique for dimension reduction (Fig. 1C).

8 major cell populations were identified, including 6 immune cell types: T cells (characterized by expression of the CD3E gene), neutrophils (S100A8), macrophage (CD68), NK cells (GNLY), B cells (MS4A1), and DC (CD1c); and 3 non-immune cell types: hepatocytes (ALB) ^24^ and endothelial cells (VWF) ^25^ (Fig. 1C, S1A). The cell counts of each population were presented in Figure. 1C and no single population was derived from one dominant sample (Fig. S1B).

To investigate the immune populations at the single-cell proteomic level, we also performed mass cytometry profiling of the same cohort. Notably, two antibody panels (lymphoid and myeloid) were created for this study based on many shared markers between scRNA-seq and CyTOF analysis (Fig 1B).

The lymphoid panel was built to quantify markers that identify lymphoid cells, such as T cell (CD3, CD4, and CD8), B cell (CD19, CD20, and CD138), and NK cell (CD56, CD57, and CD161), immunomodulatory molecules (PD-1) and chemokine reporters (CD127 and CCR7). The myeloid panel was designed to identify different populations of myeloid cells, such as macrophages (CD68, CD206, and CD86), granulocytes (CD15 and CD66b), and DCs (CD11c). Both panels included markers for the identification of immune cells (CD45) and proliferation (KI67), more details are in Figure 1D.

3,242,315 cells in total were included for Cy-TOF analysis after QC. To partition the cells into distinct phenotypes for such a high number of cells, we applied the SPADE clustering algorithm accompanied by Hierarchical Stochastic Neighbor Embedding (HSNE) to generate two-dimensional graphs within Cytosplore software ^26^. Following the scRNA-seq results, major immune cell subsets were identified (Fig. 1E and 1F, Fig. S1C and S1D).

Based on both transcriptional and proteomic profiles, we found immune cells were the majority of the BA ecosystem. More concretely, a very significant proportion of T cells, macrophages, granulocytes, and NK cells existed in the BA immune microenvironment.

### Detection of human endogenous retrovirus in BA infant family

With the evidence linking BA development to viral infection, many questions regarding the etiology of the disease remain unanswered. To elucidate the molecular and cellular mechanisms of virus-induced pathologies of BA, we mapped scRNA-seq data onto a large database of known viral genomes to systematically scan for viral RNA in host transcriptome via Viral-Track ^19^ (Fig 2A), a robust and unsupervised computational tool, in a direct mapping strategy. Employing Viral-Track on patient samples, we revealed the infection landscape of the virus and the interaction with the host. Two viral components, human endogenous retrovirus K113 (HERVK113, NC_022518) and Human parvovirus B19 (HP-B19, NC_000883), were successfully identified. 12 samples were divided into 4 subsets: 4 samples infected with HERV-K113, 1 sample with HP-B19, 1 sample detected with both HERV-K113 and HP-B19 simultaneously, the remaining 7 samples failed to be detected with any virus (Fig 2B), which indicated HERV-K113 holds the dominant position in virus coverage analysis.

**Figure 2.**
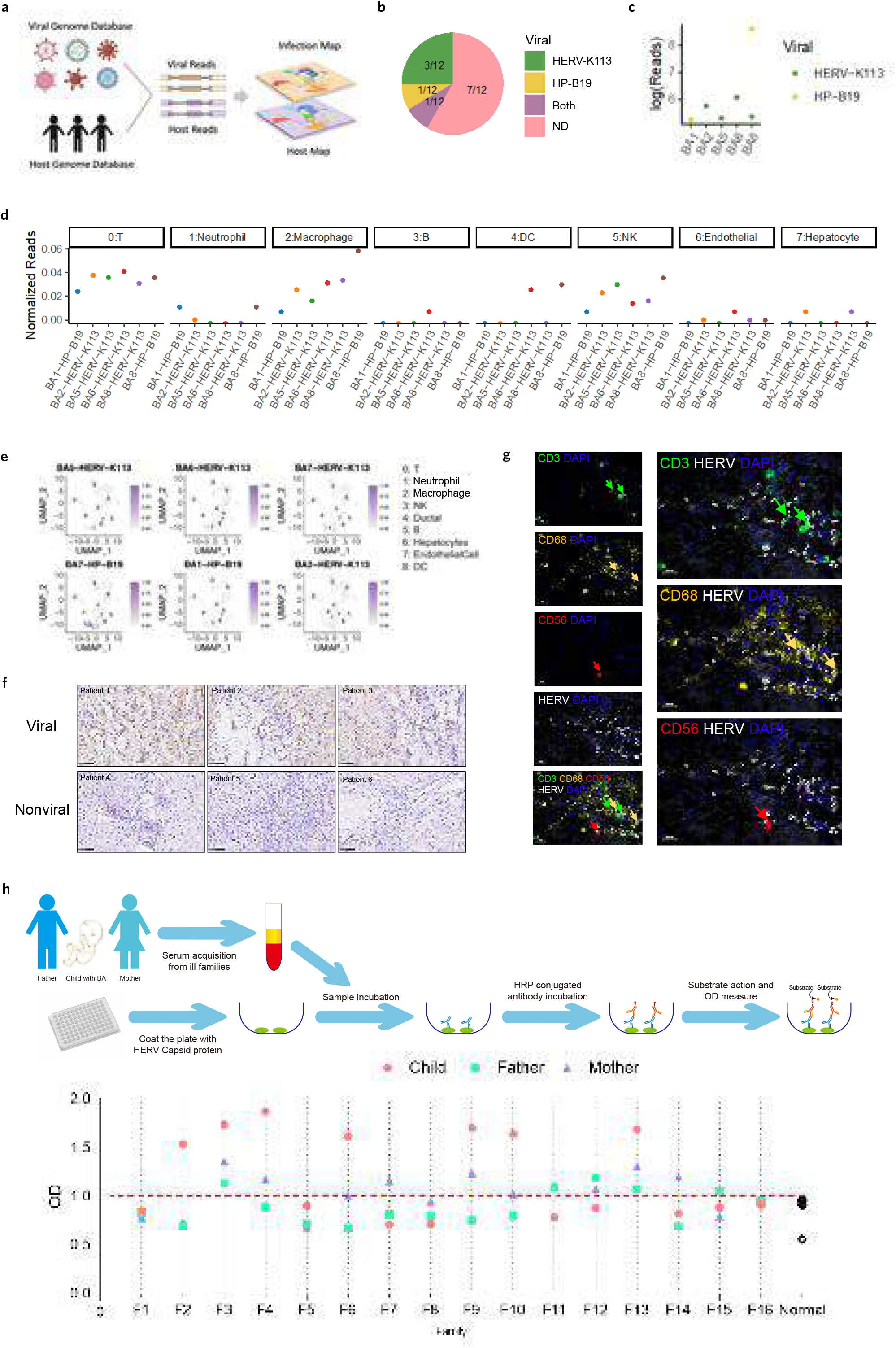
Virus detection in BA patients. A. Viral-Track algorithm for scRNA-seq data from BA patients’ samples. scRNA-seq reads were mapped to host genome and viral genome databases to define the host and virus scRNA-seq map separately. B. Pie chart showing the infection map of BA patients. HERV-K113 was detected in 3 out of 12 BA patients (green). HP-B19 virus was detected in 1 out of 12 patients (yellow). And 1 out of 12 patients was infected by both viruses (purple). Viral-Track didn’t spot any virus infection in the remaining 7 patients (pink). C. Viral reads in infected patients. Viral reads shown in log form were calculated for all the virus-infected BA patients. The blue color indicated the HERV-K113 virus and the yellow color indicated the HP-B19 virus. D and E. Viral read across major cell populations. Viral reads normalized to cell population reads were shown by dot plot and UMAP in D and E. F. IHC staining was performed to confirm the HERV existence in different BA patients. G. Multiplex IHC staining was performed to prove the HERV infection target cell in BA patients. Green, yellow, red, and white arrows indicated T cell, macrophage, NK cell, and HERV virus, respectively. And DAPI staining for the cell nucleus was shown in blue color.

HERV-K113 is the youngest and most active member of the human endogenous retroviruses of all human endogenous retroviruses known today, which is the only one that has been shown to produce viral particles ^27^. Furthermore, the minor virus component identified, HP-B19, proved the application of the Viral-Track pipeline in the BA scRNA-seq dataset was specific and successful since HP virus has been detected in the liver of BA patients ^28^. However, we mainly focus on HERV in this study, since the HP virus was established to be a noncausal agent of BA pathologies ^28^. In addition, the distribution of detected viral reads was distinct for HERV and HP virus, compared to the bi-polarized distribution mode of HP-B19 reads in 2 samples, HERV-K113 remained stable among different HERV carrying patients (Fig 2C).

To further validate the specificity of the virus detected in BA, we performed Viral-Track for 6 HCC scRNA-seq samples. 12 detailed cell clusters were found (Fig. S2A) and only HBV virus was detected in 4 out of the 6 samples (Fig. S2B). Neither HERV nor HP were found in the HCC samples.

To gain deeper insights into the distribution of infected cells in BA, we further analysed the enrichment of the infected cells. We observed a strong enrichment of viral reads in the liver resident immune cells, including T cells, macrophages, and NK cells (Fig 2D and E).

Subsequently, immunohistochemistry (IHC) against HERV capsid protein was performed to confirm the HERV activation in the BA patients. BA patients were separated into the HERV^+^ and HERV^-^ groups based on the Viral-Track results and HERV was detected in samples from the HERV^+^ group only. On the contrary, there was no signal of HERV observed in the HERV-group (Fig 2F).

Furthermore, the distribution of HERV in types of cells was verified by multiple immunohistochemistry (mIHC). Major immune residents such as T cells (identified by CD3), macrophages (identified by CD68), and NK cells (identified by CD56) were positive with HERV (Fig. 2G), shown by mIHC.

To further validate the HERV discovery in BA patients, we developed the ELISA assay for HERV-K113 capsid protein reactive antibodies (Fig. 2H). We established an independent patient cohort with 16 patients and their parents (Table S1). We found 7 out of 16 patients shows reactive antibodies to HERV-K113. Furthermore, within the 7 families, at least one of the parents also displayed reactive antibodies, and usually mother, against HERV-K113 (Fig. 2H). This result suggested HERV activation was likely to be inherited from parents.

Our results suggested a significant proportion of BA patients carrying HERV-K113 possibly through family heritage and HERV activation was mainly found in the immune residents including T cells, NK cells, and macrophages. Thus, T cells, NK cells, and macrophages were dissected in detail.

### HERV activation induces TNF producing Tem cell phenotype

To understand the immunopathogenesis mechanism related to HERV activation in T cells of BA, we applied tensor analysis for both scRNA-seq and Cy-TOF datasets ^29^. First, we built the tensor-based on scRNA-seq and Cy-TOF datasets, respectively. The sample information, cell subset clustering information, and gene expression (scRNA-seq) or protein marker expression (Cy-TOF) information, formed a 3D “tensor”. Then the tensors were decomposed and linked to virus-associated features via independent components analysis (ICA) (Fig.3A)

**Figure 3.**
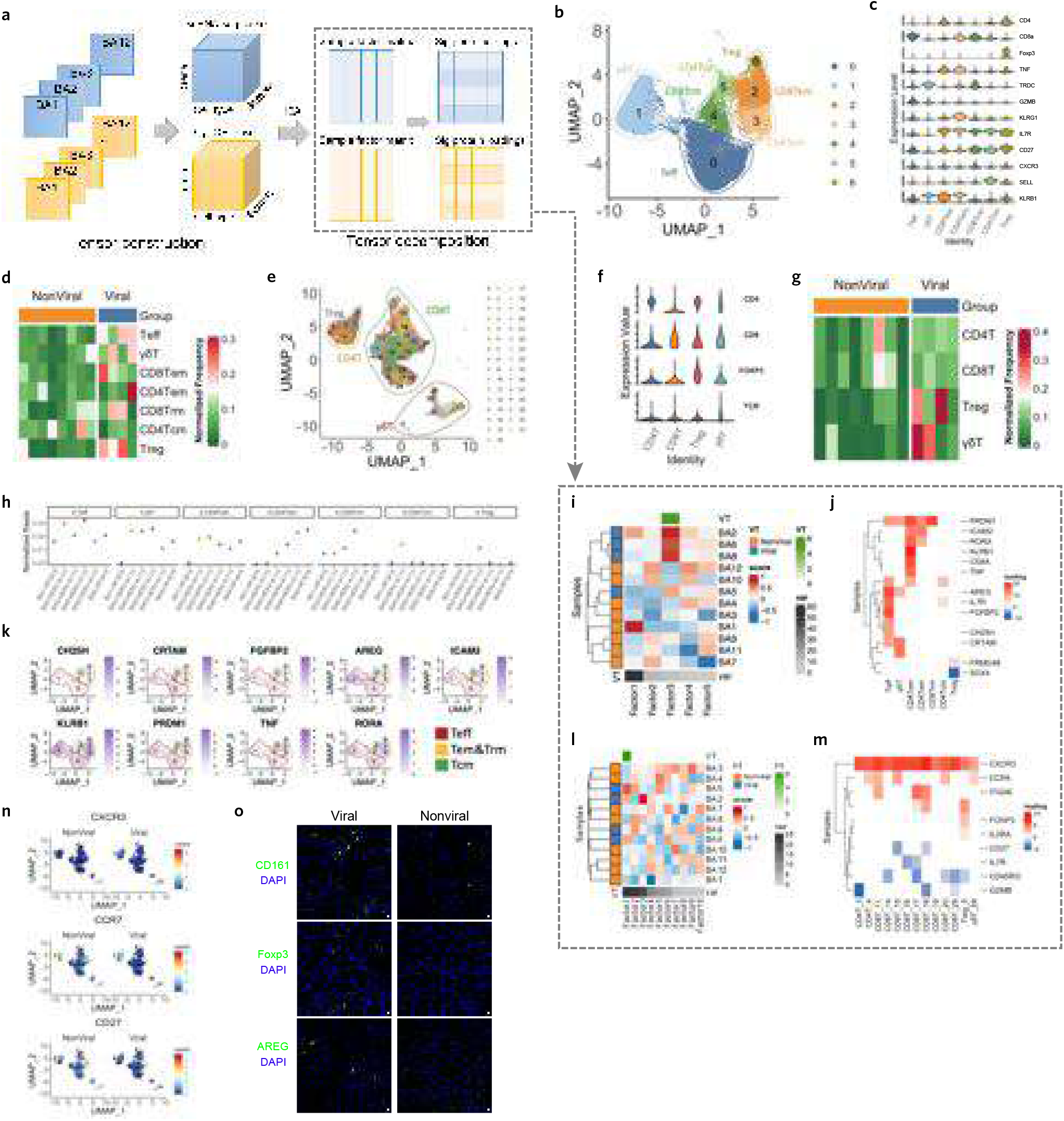
T cell re-education in BA patients. A. Tensor analysis pipeline. Two 3D “tensor” were constructed by sample, cell type, and gene expression (scRNA-seq) or protein marker expression (Cy-TOF), respectively. Then the tensors were decomposed and linked to HERV activation status via independent components analysis (ICA). B. 7 refined T cell clusters were shown by UMAP based on scRNA-seq data. C. The feature genes were shown for all the refined T cell clusters. D. The normalized frequency of refined T cell clusters calculated from 12 scRNA-seq BA samples was shown, split by the group. Orange refers to HERV^-^ NonViral, blue refers to HERV^+^ Viral. E. UMAP analysis of T cells based on the Cy-TOF results. 31 T cell clusters were found via PhenoGraph, and further annotated into 4 subpopulations. F. The typical markers were shown for Cy-TOF clusters annotation. G. The normalized frequency of refined T cell clusters calculated from 12 Cy-TOF BA samples was shown, split by the group. Orange refers to HERV^-^ NonViral group, blue refers to HERV^+^ Viral group. H. Viral reads (normalized to subset gene reads) in different T cell clusters were shown for different patients (different colours). I. The sample score matrix showed the level of each sample in present factors based on scRNA-seq data. Left annotation showed sample group information. The bottom annotation displays the variation of each factor. Top annotation was the association between sample score and group variable, shown in -log10(p-value) format. J. The corresponding loadings matric of a significant factor in Fig 3I for determining which genes from each cell type are significant. K. UMAP plots showed the expression level of selected significant genes in Fig 3J in T cells. L. The sample score matrix showed the level of each sample in present factors based on Cy-TOF data. Left annotation showed sample group information. The bottom annotation displays the variation of each factor. Top annotation was the association between sample score and group variable, shown in -log10(p-value) format. M. The corresponding loadings matric of a significant factor in Fig 3L for determining which genes from each cell type are significant. N. Group-split UMAP plots showed the expression level of significant markers in Fig 3M in T cells. O. Immunofluorescence staining of CD161, FoxP3, and AREG (green) in HERV^+^ viral and HERV^-^ non-viral patients. Blue staining of DAPI showed the cell nucleus. CD161, FoxP3, and AREG were markers for TNF^+^ CD161^+^ CD8 T cell, Treg cell, and AREG1^+^ γδ T cell, respectively.

Fine-resolution clustering of T cells revealed 7 subsets (Fig. 3B), including CD8^+^ GZMB^+^ T effector cells (cluster 0, T_eff_), TCRγ^+^ γδ T cells (cluster 1, γδT). Cluster 2 (CD8^+^) and 3 (CD4^+^) cells were IL7R^+^ KLRB^+^ KLRG^+^ CD27^-^ effector memory cells (T_em_) ^30–33^. KLRB1 (CD161) defines rapidly responding effector memory T cells ^30^ and KLRG1 also denotes effector memory T cells ^32, 33^ (Fig. 3C). While cluster 4 cells were resident CD8^+^ T cells (T_rm_) with IL7R, CXCR3 expression ^31^ (Fig.3C). And cluster 5 cells were IL7R^+^ KLRB^-^ KLRG1^-^ CD27^+^ SELL^+^ CD4^+^ central memory T (T_cm_) cells ^31, 32^ (Fig. 3C). Cluster 6 cells were FOXP3^+^ regulatory T cells (Treg) (Fig. 3C) ^35^. Please note, GZMB was mainly expressed by T_Eff_ and γδT cells while CD4 and CD8^+^ T_em_ expressed a high level of TNFα (Fig. 3C). And it seemed that T_em_ and T_rm_ cells were more enriched in HERV^+^ patients while T_cm_ cells were more enriched in the HERV^-^ patients (Fig.3D). Similarly, γδ T cells and Treg cells were also enriched more in the HERV^+^ patients.

We also performed single-cell proteomic analysis for the same cohort of patients via Cy-TOF. The T populations identified by CyTOF were clustered into 31 subsets using the PhenoGraph algorithm and embedded in two dimensions using UMAP (Fig. 3E). 4 major T cell subsets were annotated based on the protein markers, including CD4^+^ T, CD8^+^ T, Treg, and γδ T cells (Fig. 3E and F). Consistent with the scRNA-seq results, more Treg and γδ T cells were found in the HERV^+^ patients (Fig. 3G).

In addition, we observed that the majority of the viral reads were detected within the T_eff_, γδ T cells, effector memory, and resident memory T cell subsets, but not in central memory T cells or Tregs (Fig 3H), indicating that HERV activation might block the development of effector T cells into central memory phenotype.

To delineate the feature changes of the T cell lineage associated with HERV activation status, we decomposed the scRNA-seq and CyTOF tensor and linked the decomposed factor to the viral activation status via ICA, respectively (Fig. 3A). We divided 12 samples into the viral and non-viral groups with a ratio of 4:8 based on the detection of the HERV virus (Fig. 2B). After tensor decomposition, we found scRNA-seq factor 3 was correlated with viral activation status significantly (Fig. 3I). And a series of genes in various T cell subsets were identified to be associated with HERV activation (Fig. 3J).

Cholesterol 25-hydroxylase (CH25H) catalyzes the synthesis of 25HC from cholesterol. CH25H and 25HC are implicated in modulating anti-viral immunity at multiple levels ^34, 35^. CH25H was found to be more expressed in T_eff_ cells (Fig. 3K). CRTAM was found to regulate late phase T cell polarity and effector function ^36^. And we observed CRTAM enriched expression in T_eff_ and γδ T cells (Fig. 3K). FGFBP2^+^ was found to be an activation marker for effector T cells after SARS-CoV-2 infection ^37^ and we also found T_eff_ cells expressed this marker as well (Fig. 3K). All the effector T cell markers indicated their important roles in anti-HERV activation. Amphiregulin (AREG) enhances regulatory T cell-suppressive function via the epidermal growth factor receptor ^38^ and we found ARGE was highly expressed in the γδ T cells (Fig. 3K), consistent with the increased frequency of Treg in HERV^+^ BA infants (Fig. 3D), although HERV activation in Treg cells was not found. And these results suggested increased Treg frequency was an indirect effect of HERV activation in γδ T cells. ICAM2 was found to be expressed by effector memory cells (Fig. 3K) and involved in transendothelial migration of human effector memory T cells ^39^. KLRB1 (CD161) defines rapidly effector T cells associated with improved survival in HPV16-associated tumors ^30^ and we observed KLRB1 was highly expressed in Tem cells (Fig. 3K). PRDM1 (BLIMP1) enhanced the formation of effector memory CD8^+^ T cells during lymphocytic choriomeningitis virus (LCMV) infection, and PRDM1deficiency promoted the acquisition of central memory cell properties. We found PRDM1 was enriched in T_eff_ and CD161^+^ Tem cells (Fig. 3K). HERV associated upregulation of PRDM1 in effector memory cells might have blocked their development into central memory cells. We also noticed both CD4^+^ and CD8^+^ Tem cells expressed high levels of TNFα (Fig. 3K). In addition, RORA was found to be highly expressed in Tem (Fig. 3K), which promotes memory T cell formation and response to infection ^40^. All these results indicated that T cells sustained Tem state after HERV activation for better anti-viral functions.

Similarly, we performed tensor analysis for the Cy-TOF data. We found factor 1 of Cy-TOF tensor was correlated with viral activation status significantly (Fig. 3L) and a series of protein markers were revealed by detailed analysis of factor 1 (Fig. 3M). Among them, CXCR3 (Trm marker) was more expressed in viral patients while CCR7 and CD27 (Tcm markers) were more expressed in non-viral patients (Fig. 3N), consistent with scRNA-seq results.

Furthermore, scRNA-seq and Cy-TOF tensor analysis results were further validated by immunofluorescent staining. CD161^+^ Tem cells were more frequently detected in the viral group of patients. And AREG+γδ T and Treg cells were more frequently detected in the viral group of patients (Fig.3O).

Collectively, these data highlighted that T cell subsets shifted more to the anti-viral effector response after HERV activation, such as increase of CH25H in Teff cells, increased frequency of effector T cells, and CD161^+^ TNFα^+^ Tem cells. In addition, Tregs were also increased in the viral group of patients, possibly due to enhanced AREG signal via γδ T cells.

### HERV activation leads to distinct licensing factor engagement of NK cells

We next interrogated the HERV associated features of NK cells in BA samples. NK cell subsets annotations were refined further on basis of differences in the expression of specific markers. 8 rarefied subpopulations were distinguished (Fig 4A-B), including CX_3_CR1^+^ NK ^41^, FOS^+^ NK ^42^, KIR2DL1^+^ NK, PTGDS^+^ NK, KLRC^+^ NK ^43, 44^, RAMP^+^ NK ^45^ and CCL3^+^ NK ^46^.

**Figure 4.**
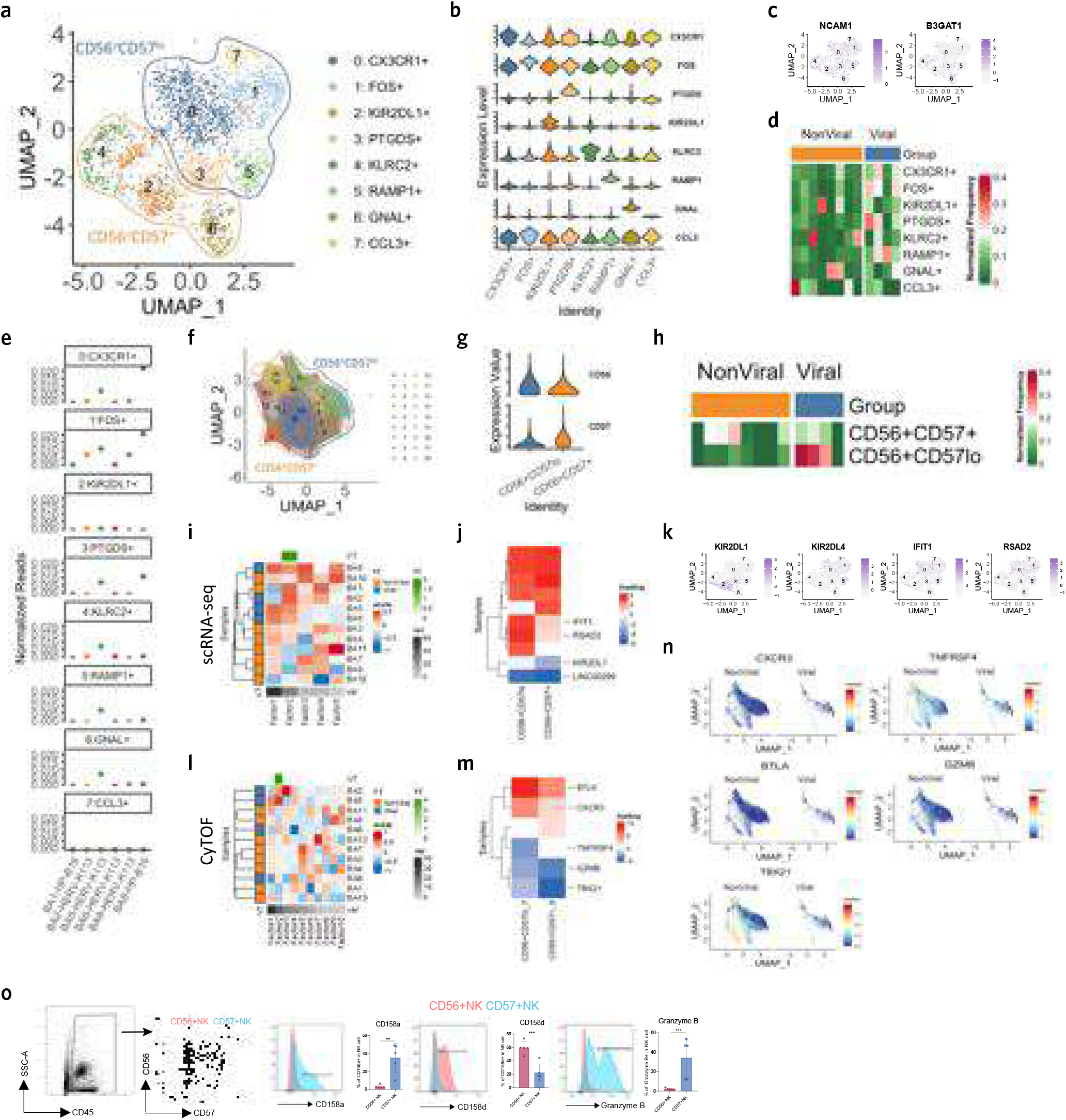
Distinguished licensing factor engagement of NK cell in the viral and non-viral groups of BA patients. A. 8 (0-7) refined NK cell clusters were shown by UMAP based on scRNA-seq data. B. The specific feature genes were shown for all the refined NK cell clusters. C. The expression pattern of NCAM1 (CD56) and B3GAT1 (CD57) were shown in UMAP. D. The normalized frequency of refined NK cell clusters calculated from 12 scRNA-seq BA samples was shown, split by the group. Orange refers to HERV^-^ NonViral group, blue refers to HERV^+^ Viral group. E. Viral reads (normalized to cell subset gene reads) in different NK cell clusters were shown for different patients (different colours). F. UMAP analysis of NK cells based on the Cy-TOF results. 18 NK cell clusters were found via PhenoGraph and further divided into CD56^+^ CD57^+^ and CD56^+^CD57^low^. G. Violin plot displayed the expression pattern of NCAM1 (CD56) and B3GAT1 (CD57) on Cy-TOF data. H. The normalized frequency of refined NK cell clusters calculated from 12 Cy-TOF BA samples was shown, split by the group. Orange refers to HERV^-^ NonViral group, blue refers to HERV^+^ Viral group. I. The sample score matrix showed the level of each sample in present factors based on scRNA-seq data. Left annotation showed sample group information. The bottom annotation displays the variation of each factor. Top annotation was the association between sample score and group variable, shown in -log10(p-value) format. J. The corresponding loadings matric of a significant factor in Fig 4I for determining which genes from each subpopulation are significant. K. UMAP plots showed the expression level of significant genes in Fig 4J in NK cells. L. The sample score matrix showed the level of each sample in present factors based on Cy-TOF data. Left annotation showed sample group information. The bottom annotation displays the variation of each factor. Top annotation was the association between sample score and group variable, shown in -log10(p-value) format. M. The corresponding loadings matric of a significant factor in Fig 4L for determining which genes from each subpopulation are significant. N. Group-split UMAP plots showed the expression level of significant markers in Fig 4M in NK cells. O. FACS results for CD56+ and CD57+ NK cell clusters. A brief gating strategy was shown in the left panels. CD56+ and CD57+ NK cells were gated after CD45+ immune cell gating. Representative histogram and statistical plots showed the comparison of expression level for CD158a, CD158d, and Granzyme B between CD56+ and CD57+ NK cells (** p<0.01, ***p<0.001).

NK cell subsets could be categorized into two major compartments, CD56^+^ CD57^+^ and CD56^+^CD57^low^ (Fig. 4A and C). And CD56^+^CD57^low^ NK cells (cluster 0, 1, 3, and 5), showed an increased fraction in the viral group of patients (Fig. 4D). Furthermore, we found CD56^+^CD57^low^ NK cells (clusters 0, 1, 3, and 5) were primary targets of HERV activation (Fig. 4E).

To further depict the importance of NK subpopulations in viral-induced immune response, we analysed the variety of subpopulation fractions between viral and non-viral groups based on Cy-TOF data as well. We discovered 18 clusters embedded in the two-dimensional UMAP diagram via the PhenoGraph re-cluster (Fig. 4F). Consistent with the scRNA-seq results, 18 Cy-TOF NK clusters comprised two primary categories, CD56^+^ CD57^+^ and CD56^+^CD57^low^ cells (Fig. 4G). In addition, we also observed higher CD56^+^CD57^low^ clusters in the viral group of patients via Cy-TOF (Fig. 4H).

Next, we wanted to illustrate the mechanism of the imbalanced NK cell distribution after HERV activation. Tensor analysis was formed for NK cells based on scRNA-seq and Cy-TOF data. We found factor 2 of the scRNA-seq tensor was significantly associated with the HERV activation (Fig. 4I). And further dissection of factor 2 revealed that CD56^+^ CD57^low^ NK cells expressed a higher level of IFIT1 and RSAD2 (Fig. 4J-K). Both IFIT1 and RSAD2 are important for NK antiviral functions ^47–51^.

NK cell killing function depends on proper licensing molecule engagement ^55^, such as killer cell immunoglobulin-like receptor (KIR). Based on the scRNA-seq tensor analysis, we found the two KIRs were distinguished in CD56^+^ CD57^+^ and CD56^+^CD57^low^ NK cells (Fig. 4J-K).

CD158a (KIR2DL1) were mainly found in the CD56^+^ CD57^+^ NK subsets while CD158d (KIR2DL4) were mainly found in the CD56^+^CD57^low^ NK subsets (Fig. 4K).

In addition, we also performed tensor analysis for the Cy-TOF data and we found Cy-TOF tensor factor 2 was positively associated with the HERV activation in NK cells (Fig. 4L). And detailed analysis for factor 2 showed higher expression of BTLA and CXCR3 in the CD56^+^ CD57^low^ NK cells (Fig. 4M-N). Both BTLA and CXCR3 were shown to induce immunosuppressive NK phenotypes ^52, 53^. In contrast, TNFRSF4 (CD134), GZMB, and TBX21 (TBET) were higher in the CD56^+^ CD57^+^ cells (Fig. 4M-N). All these 3 markers were important for the cytotoxic activities of NK cells ^54, 55^.

To further confirm our results based on scRNA-seq and Cy-TOF. Classical flow cytometry was performed to verify that CD56^+^CD57^low^ NK cells expressed a higher level of CD158d. On the contrary, another type of NK cells identified by CD56^+^ CD57^+^ was detected with higher CD158a-expression (Fig. 4O). Furthermore, CD56^+^ CD57^+^ NK cells showed a higher level of Granzyme B than CD56^+^ CD57^low^ NK cells, consistent with the CyTOF results.

Collectively, the results of this exploratory scRNA-seq and Cy-TOF experiments revealed the presence of two phenotypically distinct NK cells with different immune statuses in patients with BA, and the difference of cytotoxicity was a consequence of distinct licensing factor engagement in NK cells associated with HERV activation.

### HERV activation is mainly detected in the chemokine deficient FOLR2^+^ resident macrophage

Considering macrophages comprised a significant portion of infected cells in BA infants, we next inspected macrophage-specific features associated with HERV activation at refined resolution.

Macrophages could be further divided into 7 subsets (Fig 5A-B), including S100A9^+^ macrophage (cluster 0) ^56^, HLA II^+^ macrophage (cluster 1), CD5L+ macrophage (cluster 2) ^57^, KLF2^+^ macrophage (cluster 3) ^58^, GPR183^+^ macrophage (cluster 4) ^59^, MT1G^+^ macrophage (cluster 5) ^60, 61^ and CCL8^+^ macrophage (cluster 6) ^62^. We observed the 7 macrophage subsets harboured two major categories, resident (cluster 1, 2 and 5) and infiltrating macrophages (cluster 0, 3, 4 and 6) (Fig. 5A).

**Figure 5.**
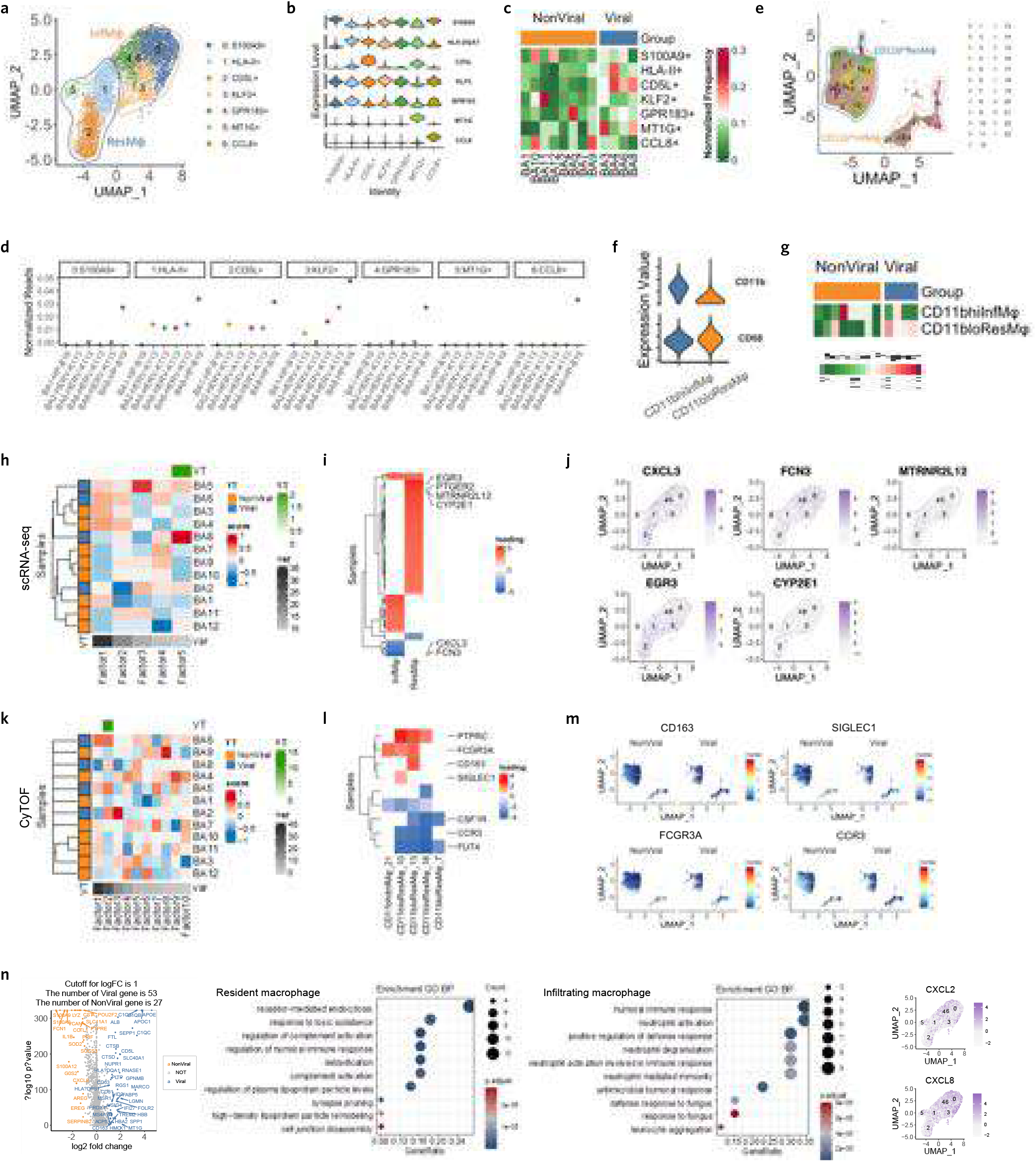
Macrophage function shift in the viral and non-viral groups of BA patients. A. 7 (0-6) refined macrophage cell clusters were shown by UMAP based on scRNA-seq data. The UMAP plot showed the source of the macrophage clusters. The blue colour indicated resident macrophages and the yellow colour indicated infiltrating macrophages. B. The specific feature genes were shown for all the refined macrophage cell clusters. C. The normalized frequency of refined macrophage cell clusters calculated from 12 scRNA-seq BA samples was shown, split by the group. Orange refers to NonViral, blue refers to Viral. D. Viral reads (normalized to gene reads of macrophage clusters) in different NK cell clusters were shown for different patients (different colours). E. UMAP analysis of macrophage cells based on the Cy-TOF results. 23 macrophage cell clusters were found via PhenoGraph and then divided into 2 subpopulations. F. Violin plots showed the expression level of CD11b and CD68 in macrophages. G. The normalized frequency of refined macrophage cell clusters calculated from 12 Cy-TOF BA samples was shown, split by the group. Orange refers to HERV^-^ NonViral, blue refers to HERV^+^ Viral. H. The sample score matrix showed the level of each sample in present factors based on scRNA-seq data. Left annotation showed sample group information. The bottom annotation displays the variation of each factor. Top annotation was the association between sample score and group variable, shown in -log10(p-value) format. I. The corresponding loadings matric of a significant factor in Fig 5I for determining which genes from each subpopulation are significant. J. UMAP plots showed the expression level of significant genes in Fig 5J in macrophages. K. The sample score matrix showed the level of each sample in present factors based on Cy-TOF data. Left annotation showed sample group information. The bottom annotation displays the variation of each factor. Top annotation was the association between sample score and group variable, shown in -log10(p-value) format. L. The corresponding loadings matric of a significant factor in Fig 5L for determining which genes from each subpopulation are significant. M. Group-split UMAP plots showed the expression level of significant markers in Fig 5M in macrophages. N. The volcano plot showed the DEGs between resident macrophages (blue colour) and infiltrating macrophages. And GO analysis was performed for the DEGs.

Resident macrophage was characterized by the broad expression of genes reported in tissue-resident macrophages, such as complement proteins (C1QA and C1QB) ^58, 59^, folate receptor (FOLR2) ^60,61^and scavenger receptor (CD163, MRC1, and MARCO) ^62^ (Fig S3A). And FOLR2 is also recognized as the marker for fetal-derived tissue-resident macrophage in the liver ^60, 61^.

The inflammatory markers, IL1B, CXCL2, CXCL8, CD11b (encoded by ITGAM), and CCR2 were centralized in the infiltrating macrophage (Fig S3A). Consistent with our previous findings that liver infiltrating macrophages derived from bone marrow were mainly CD11b^+^ in both human and mouse ^63, 64^. In addition, we noticed resident macrophages were more enriched in the HERV^+^ patients while infiltrating macrophages were more enriched in the HERV^-^ patients (Fig. 5C).

Previously, we have shown that liver resident macrophages were derived from fetal definitive haematopoiesis ^63–66^ based on fate-mapping results. Single-cell lineage tracing analysis based on mitochondria ^67^ and genomic mutations showed that resident macrophages carried unique mutations derived from early fetal definitive haematopoiesis. And a big portion of the mutations was shared between infiltrating and resident macrophages, derived from the major definitive haematopoiesis wave (Fig.S3B and C), consistent with our previous fate-mapping results. Viral-Track results showed that the fetal-derived tissue-resident macrophage (cluster 1 and 2) were the major targets for HERV activation ^60^ and infiltrating macrophage was less susceptible to HERV activation (Fig. 5D).

In addition, we also performed an in-depth phenotypical analysis of macrophage cells by mass cytometry (Fig 5E), a total of 23 macrophage clusters were identified (Fig 5E). Consistent with the scRNA-seq data, the 23 clusters could be categorized into CD11b^+^ CD68^+^ infiltrating macrophages and CD11^low^ CD68^+^ residents (Fig. 5E and F). In addition, Cy-TOF data also proved that resident macrophages were more enriched in the HERV^+^ patients (Fig. 5G).

To unravel the HERV associated features of macrophages, tensor analysis was also formed for both scRNA-seq and Cy-TOF data. Factor 5 of the scRNA-seq tensor was positively correlated with the HERV activation and detailed analysis of factor 5 revealed several feature genes related to HERV activation in macrophage (Fig. 5H and I). CXCL3, FCN3 enrichment was associated with HERV^-^ status of infiltrating macrophages (Fig. 5J). CXCL3 might contribute to the neutrophil accumulation in the HERV^-^ BA patients based on the CXCL3/CXCR2 neutrophilic chemotactic axis ^68^. Ficolin-3 (FCN3) level was reported to be lowered in HIV patients and its close homolog, FCN2 was previously shown to confer resistance to several hepatotropic viruses ^69^. It was likely that FCN3 upregulation in infiltrating macrophages confers HERV resistance as well. Similarly, MTRNR2L12, EGR3, and CYP2E1 enrichment were associated with the HERV^+^ status of resident macrophages (Fig. 5I-J). MTRNR2L12, an anti-apoptotic lncRNA, was significantly upregulated in the severe COVID-19 patients ^70^.

MTRNR2L12 expression might contribute to the sustaining of resident macrophages after HERV activation. ERG3 was reported to be associated with HIV reactivation ^71^, consistent with its expression in resident macrophages. And CYP2E1 was reported to attenuate the interferon antiviral response ^72^, indicating the possible mechanism of HERV susceptivity of resident macrophage.

We also performed tensor analysis for the Cy-TOF data and factor 2 of the tensor was positively correlated with the HERV activation (Fig. 5K). CD163, FCGR3A, and SIGLEC-1 enrichment were associated with the HERV^+^ status of resident macrophages (Fig. 5L and M). These markers were all specific to resident macrophages ^65, 73, 74^ and associated with retrovirus infection. While CCR3 downregulation was associated with the HERV^+^ status of resident macrophages (Fig. 5L and M), consistent with the neutrophil depletion in the HERV^+^ BA patients.

Next, we wanted to explore the differences between resident and infiltrating macrophages. The DEGs were generated via comparison of resident and infiltrating macrophages and GO analysis was also performed (Fig. 5N). We noticed resident macrophages were more specialized in the detoxification process while infiltrating macrophages was a strong neutrophil attractor and activator (Fig. 5N). For example, the higher levels of expression of CXCL2 and CXCL8 that attract and activate neutrophils ^75^, was noticed in the infiltrating macrophages (Fig. 5N), consistent with the tensor analysis.

Transcriptomic and proteomic profiling identified the resident and infiltrating macrophages. HERV activation was mainly detected in the FOLR2^+^ fetal-derived resident macrophages and associated with the increased frequency of resident macrophages. HERV activation within resident macrophages was associated with neutrophil recruitment chemokines deficiency.

### SOD1/2^+^ neutrophils mediated ROS scavenging is impaired in BA

In response to many types of infection, neutrophils are thought to take on a pivotal role in shaping the immune response. As described above, HERV activation could contribute to the remodelling of neutrophils driven by macrophages. To extend our knowledge on the role of neutrophils in HERV^+^ BA, we conducted scRNA-seq experiments relying on UMAP analysis.

The delicate clustering distinguished three distinct subpopulations (Fig. 6A-B), characterized by chemokines (CCL4L2, CXCL2, CCL3, and CCL4), calprotectin (S100A12, S100A8, and S100A9), and cytotoxic agents associated genes (NKG7, AREG, GZMB, and GNLY). The GZMB^+^ neutrophil was also identified in the liver by us in a previous study ^63^. The top 10 cluster-specific features were presented in Fig 6B.

**Figure 6.**
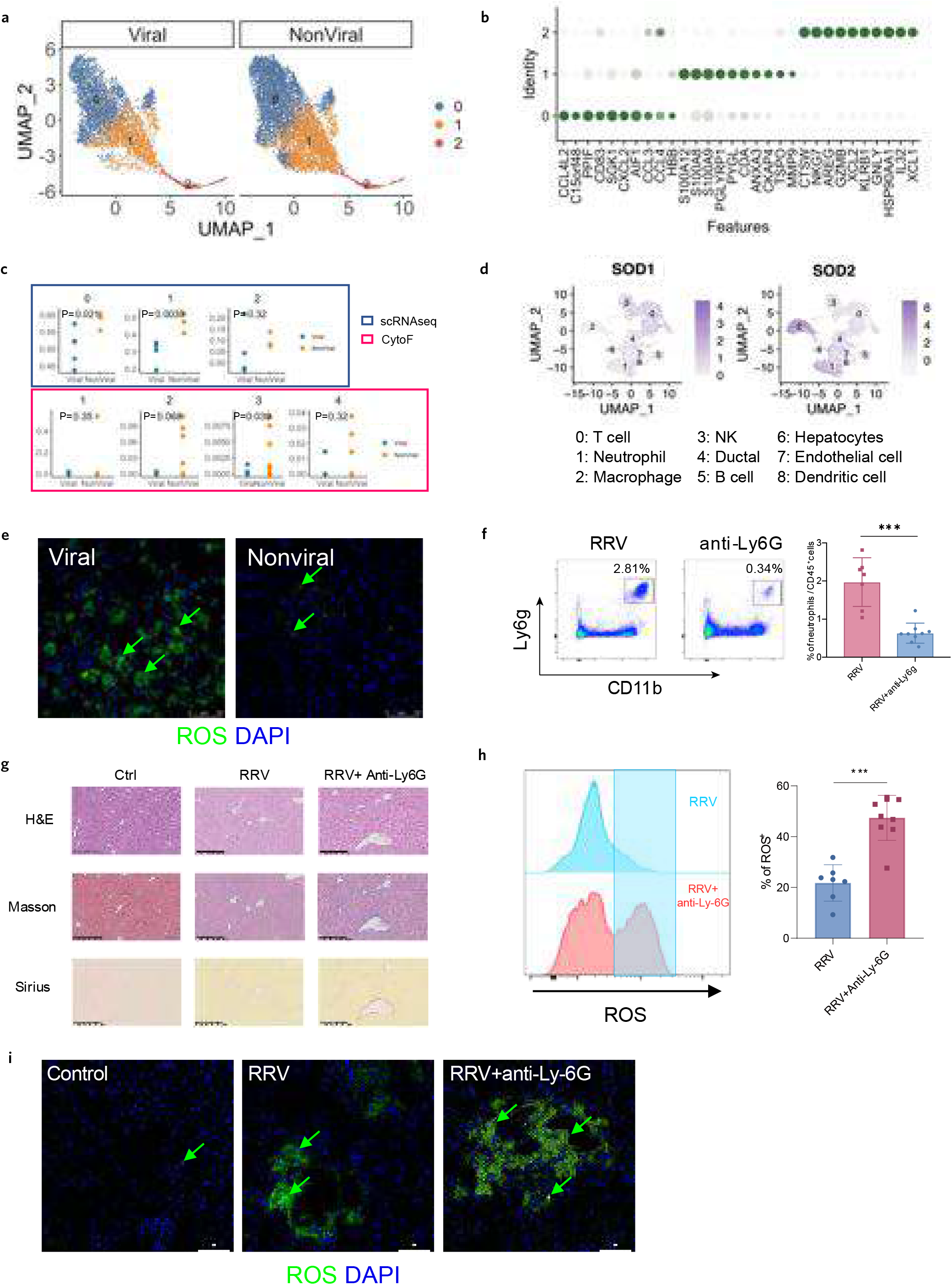
ROS scavenging neutrophils diminished in the viral group of BA patients. A. The UMAP plot showed the distribution of the 3 neutrophil clusters in the HERV^+^ viral and HERV^-^ nonviral group of BA patients. B. The top 10 feature genes were shown for all the refined neutrophil cell clusters. C. Neutrophil cell cluster frequency comparison between HERV^+^ viral (blue circle) and HERV^-^ non-viral (yellow circle) group of patients. The result in the upper panel was generated from scRNA-seq data and the result in the lower panel showed the comparison of neutrophil cell clusters from Cy-TOF data. P-value was shown in the middle of each plot. D. UMAP plots showed the expression level of SOD1 and SOD2 in neutrophils. E. Immunofluorescence staining of ROS probe (green) in HERV^+^ viral and HERV^-^ non-viral patients. Blue staining of DAPI showed the cell nucleus. F. FACS and statistical results for neutrophil depletion were shown lower panel. Neutrophil was gated as CD11b+ Ly6g+ cells after CD45+ immune cell gating (not shown, ** p<0.01, ***p<0.001).s G. HE staining showed the tissue histology. Masson and Sirus staining showed the fibrosis extent of the livers from the ctrl mice, RRV infected mice, and RRV infected mice with neutrophil depletion. H. Representative FACS histogram and statistical plots showed the comparison of ROS probe level of the livers from the RRV infected mice and RRV infected mice with neutrophil depletion. I. Immunofluorescence staining of ROS probe (green) in livers from RRV infected mice and RRV infected mice. Blue staining of DAPI showed the cell nucleus.

Strikingly, enormous neutrophils were accumulated in patients without viral infection (Fig 6A). The fraction of all 3 subpopulations of neutrophils in patients with the virus was dramatically lower than the corresponding control, which was concordant with the marked imbalance distribution in UMAP (Fig 6A).

The findings were further confirmed by CyTOF data. The associated CyTOF neutrophil clusters (clusters 1,2,3 and 4 of neutrophil) demonstrated consistent accumulation trend in HERV^-^ nonviral group of patient samples (Fig 6C).

Furthermore, we found the members of the iron/manganese superoxide dismutase family, SOD1 and SOD2, were concentrated in neutrophils, especially for SOD2 (Fig 6D), indicating the release of reactive oxygen species (ROS), which could be cleared more efficiently in the nonviral group (Fig. 6E) through ROS scavenging neutrophils. Neutrophil shortage might contribute to cell damage and death in HERV carrying patients ^76^.

To validate the ROS protection role of SOD1/2^+^ neutrophils in BA, we established a mouse BA model via Rhesus Rotavirus (RRV) induction (Fig. S4) to verify the ROS protection capacity of neutrophils. Mice injected with Rhesus Rotavirus (RRV) were administrated with anti-Ly-6G monoclonal antibody (Fig 6F). After the administration of Ly-6G mAb, neutrophils were reduced significantly (Fig. 6F). And the livers presented more serious fibrosis (Fig 6G). Meanwhile, the ROS level also increased significantly compared to the RRV group via FACS and IF (Fig 6H and I).

Thus, HERV re-education of immune residents in BA patients also greatly reduced the short-lived SOD1/2^+^ neutrophil, thus exacerbating the tissue damage caused by ROS.

### Macrophage modulation therapy for BA

Next, we planned to modulate the ecosystem of BA to alleviate liver tissue damage in BA patients.

A novel Folr2-eGFP-DTR mouse strain was developed to deplete the HERV susceptible FOLR2^+^ macrophages (Fig. 7A). BA model induced by RRV was established in the Folr2eGFP-DTR mouse (Fig. 7A) and clear ablation of FOLR2^+^ (eGFP^+^) macrophage could be observed (Fig. 7B). Meanwhile, neutrophils were increased significantly (Fig. 7C). As a result, neutrophil-mediated ROS protection was enhanced in the FOLR2^+^ macrophage depletion model and ROS level was also reduced in the RRV mediated BA model (Fig. 7D). Significantly reduced fibrosis was observed in (Fig. 7E).

**Figure 7.**
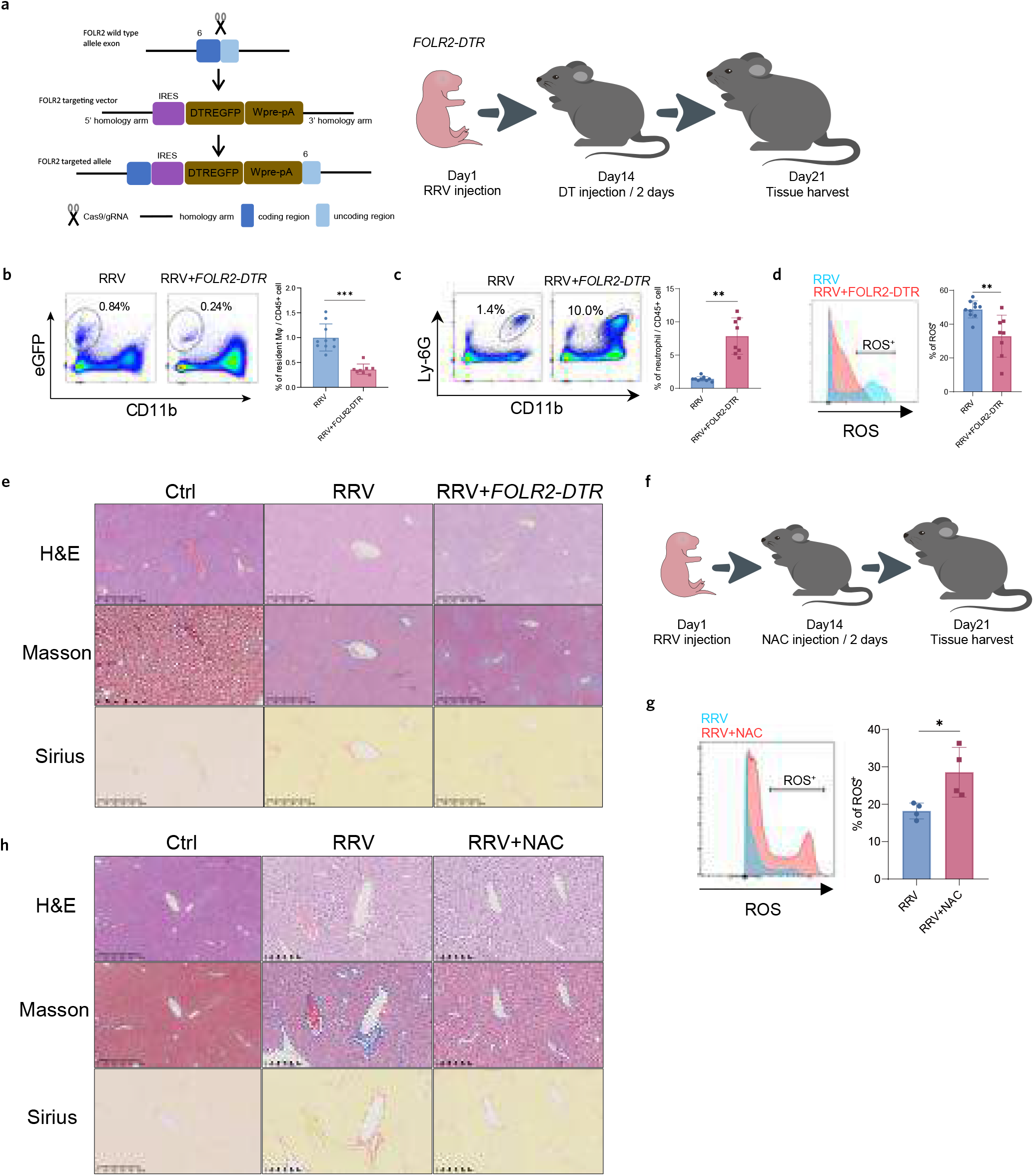
Potential BA treatment strategies. A. The construction map of the FOLR2-DTR mouse is shown in the left panel. The IRES-DTR-EGFP fusion gene was inserted into the 3’-UTR region of the FOLR2 gene locus. The mouse experiment schedule is shown in the right panel. BA was induced in FOLR2-DTR mice by RRV infection and FOLR2^+^ macrophage was depleted by DT injection. B. Folr2^+^ macrophage depletion results in mouse BA model were shown. FACS and statistical results for FOLR2^+^ macrophage depletion were shown. FOLR2^+^ macrophage was gated as CD11b^-^ EGFP^+^ cells after CD45^+^ immune cell gating (not shown, ** p<0.01, ***p<0.001). C. FACS and statistical results for neutrophils in mouse BA model with or without FOLR2+ macrophage depletion were shown. Neutrophil was gated as in Fig. 6 (** p<0.01, ***p<0.001). D. Representative histogram and statistical plots showed the comparison of ROS probe level of the livers from the RRV infected mice with or without FOLR2^+^ macrophage depletion (** p<0.01, ***p<0.001). E. HE staining showed the tissue histology. Masson and Sirus staining showed the fibrosis extent of the livers from the ctrl mice and RRV infected mice with or without FOLR2^+^ macrophage depletion. F. NAC treatment schedule. BA was induced in mice by RRV infection and NAC was injected every 2 days for 14 days. G. Representative histogram and statistical plots showed the comparison of ROS probe level of the livers from the RRV infected mice with or without NAC treatment (* p<0.05). H. HE staining showed the tissue histology. Masson and Sirus staining showed the fibrosis extent of the livers from the ctrl mice and RRV infected mice with or without NAC treatment.

We also searched for clinically proven drugs to scavenge ROS for BA treatment. NAC is a potent antioxidant and was first approved by the FDA as a respiratory drug in 1963. It is also used to prevent serious liver damage from acetaminophen poisoning. Thus, we tested its application in the treatment of BA. NAC was administrated into RRV induced BA model mouse (Fig. 7F). We found NAC could wipe off the ROS in the RRV induced BA model (Fig. 7G). In addition, liver fibrosis conditions were alleviated in the NAC treatment group (Fig. 7H).

Overall, we found BA could be potentially treated through specific macrophage modulation and NAC treatment.

## Discussion

In this study, we constructed a bimodal single-cell atlas of immune cells from BA patients through scRNA-Seq and CyTOF. Our tensor analysis provided several new sights into BA pathogenesis. Firstly, we detected HERV in BA patients via Viral-Track, IHC and mIHC at single-cell resolution. Importantly, HERV activation could be inherited from the parent, validated by ELISA. HERV was predominantly enriched in liver resident immune cells, including FOLR2^+^ macrophages, T cells, and NK cells. We noticed HERV infection could re-educate the immune residents. Importantly, we found SOD1/2^+^ neutrophils mediated ROS scavenging was impaired in BA due to FOLR2^+^ fetal resident macrophage re-education, and depletion of FOLR2^+^ resident macrophage could rescue the tissue damage in BA.

Our single-cell transcriptomic and proteomic profiling of BA samples reveals novel mechanisms of immune response in viral-associated pathological features, which was not demonstrated in the last comprehensive immune study of BA ^18^. Jun *et al* revealed pathogenic autoimmune B cells within BA immune profiling and proposed B cell modulation therapy for BA ^18^. Our results were consistent and complement with Jun *et al*’s findings in terms of macrophage and B cells. They noticed hepatic macrophages have decreased inflammatory function in infants with BA and we further elaborated that this phenomenon could be due to HERV activation within the resident macrophages. In addition, expansion of autoreactive B cells was detected in infants with BA and it was likely that a portion of the autoreactive B cells was in fact against the HERV as we showed here. Jun *et al* advanced the hypothesis that B-cell-modifying therapies facilitate immune recovery in BA patients. Complimentary, we highlighted the virus-associated mechanisms and showed that the modification of ROS in viral infection BA patients may ameliorate BA pathology.

Numerous studies have reported the inconclusive relationship between the virus and BA pathogenesis ^3–15^. For example, the detection of the HP virus in the liver appears to be no connection with BA ^15^. In our study, HERV was detected sporadically in human BA, which was consistent with the previous findings that autoimmunity and immune dysregulation possibly contributed to BA pathogenesis ^77^. The autoimmunity and immune dysregulation were likely induced by HERV activation in BA patients. In patients without the presence of HERV, the pathogenesis of BA may be triggered by bile duct epithelial injury caused by physical or chemical insulting.

We noticed HERV activation was more frequent in the CD56^+^ CD57^low^ NK cells and associated with decreased cytotoxicity. In addition, KIR2DL1(CD158a) was mainly detected in the CD56^+^ CD57^+^ NK cells and KIR2DL4 (CD158d) was mainly detected in the CD56^+^ CD57^low^ NK cells. Our findings seemed against the CD158a inhibition role defined previously ^78^. However, in the HIV patients, KIR2DL1/HLA–C2 interaction was reported to confer enhanced antibody-dependent cellular cytotoxicity (ADCC) in vaccine-conferred protection from infection. Among effector cells that mediate ADCC are CD57^+^ natural killer (NK) cells ^79^, in good accordance with our results.

Resident macrophages in the BA infants were carrying detoxification functions. We thought it might be due to the analysis was performed for the disease livers, which were collected at least several months after diagnosis and the jaundice was usually quite heavy during surgery time. The initial pathogenesis phase was likely in the gestation period, and we guess macrophage during the initial phase of BA might also contribute to bile ductal epithelial cell hyperplasia as well.

In summary, our results showed HERV activation could remodel the local-host environment, contributing to impairment of ROS scavenging in BA. Consequently, our findings uncover the virus-associated mechanism of BA pathogenesis, which could provide novel insights into BA and reveal the potential of macrophage-based therapy for BA. Our results indicated HERV activation in the BA infants could be inherited from one of their parents and the application of screening strategy showcased in our manuscript could contribute to future diagnosis strategy for HERV. And our study may shed light on the important roles of HERV in other autoimmune diseases and tumours.

## Materials and Methods

### Ethics approval

All animal experiments were approved by the Research ethic committee of the First Affiliated Hospital, College of Medicine, Zhejiang University, ID of the approval: 2019-1257. Studies using human specimens were approved by the Ethics Committee of the First Affiliated Hospital, Zhejiang University School of Medicine.

### Patients and samples

Liver sample tissues from 12 patients who were diagnosed with biliary atresia without Kasai portoenterostomy were isolated by professional surgeons (L.T. and X.B.) during liver transplantation. All samples were verified by pathological examinations. Blood another 16 patients

### Animal studies

C57BL/6J and BALB/c mice were purchased from the Model Animal Research Center of Nanjing University (China). *FOLR2-DTR* mice were generated by Shanghai Model Organisms Center, Inc. (Shanghai, China), as described in the results. *FOLR2-DTR* mice were backcrossed to BALB/c mice for at least 4 generations before RRV induction. All mice were housed in the SPF facility of the First Affiliated Hospital, Zhejiang University School of Medicine with approval from the Institutional Animal Care & Use Committee (IACUC).

Rhesus Rotavirus (MMU 18006) (VR-1739, ATCC) was propagated in MA104 cells overlaid with serum-free DMEM (SF DMEM)+ 4 μg/mL trypsin for 48h. Then we performed 1: 10 serial dilutions of RRV to inoculate MA104 cells. After 15h, cells were incubated with anti-rotavirus antibody (ab181695, Abcam) followed with secondary fluorescent antibody incubation (4412, Cell Signaling Technology). We calculate the number of foci with the formula: (average number of foci between duplicates × 2.8) × 100 × virus dilution = focus-forming units (FFU)/mL.

Then we injected newborn mice intraperitoneally (i.p.) at a dose of 1.25 × 10^6^ FFU of RRV diluted in saline per gram body weight within 24h of birth. Monitor pups for 21 days for weight gain along with the clinical signs of hepatobiliary injury— jaundice and acholic stools. And on day 21, the Venous blood of mice was taken from the portal vein. Serum alanine aminotransferase (ALT), aspartate aminotransferase (AST), and total bilirubin (TBIL) levels were measured by detection kits (C019-1, C009-2, and C010-2, Nanjing Jiancheng Bioengineering Insitute). Then liver tissues were acquired to be diagnosed via pathological staining.

For depletion of neutrophils, anti-mouse Ly6G (BE0075-1, BioXcell) at the dose of 10 mg/kg was applied to the 2-week BA mice 2 times a week.

For depletion of FOLR2^+^ macrophages, *FOLR2-DTR* mice infected with RRV were injected with DT (D0564, Sigma-Aldrich) at the dose of 10 μg/kg 4 times a week.

For scavenging ROS, the mice with BA were administrated with NAC (A7250, Merck) at the dose of 300 mg/kg 4 times a week.

### Immunofluorescent imaging

Six-μm frozen slides were prepared from BA samples to be fixed through chilled methanol for 10 min. After blocking with SuperBlock (37515, Thermofisher), primary antibodies CD161 (ab259916, Abcam), Foxp3 (98377, Cell Signaling Technology) or AREG (ab234750, Abcam) followed with AF488 anti-rabbit secondary antibody (4412, Cell Signaling Technology) incubation. Then, slides were stained with DAPI for 5 min at RT. All immunofluorescent images were scanned via a four-laser scanning confocal microscope (Leica TCS SP8, Germany).

### Pathological tissue imaging

Hematoxylin & eosin, Masson (G1340, Solarbio), and Sirius (G1470, Solarbio) staining were performed to verify BA pathology. IHC experiments were processed to detect the existence of HERV (LS-C65284, LSBio). In addition, traditional IHC detecting CK19 (ab52625, Abcam) was used to portray the malformation of bile ducts in RRV-infected BA mice. Images were captured by optical microscopy (Leica DM2500 LED, Germany).

### Multiplex immunohistochemistry

For identifying the location of HERV, multiplex IHC was performed. We adopted Opal Polaris^TM^ 7-color Manual IHC Kit (NEL861001KT, Akoya). Primary antibodies including HERV (LS-C65284, LSBio), CD3 (85061S, Cell Signaling Technology), CD68 (ab213363, Abcam), CD56 (ab237708, Abcam) were involved in the mIHC. After DAPI staining, visualization of 4-color opal slides was performed by Vectra Polaris Quantitative Pathology Imaging Systems (Akoya, USA).

### ELISA

An uncoated ELISA plate (423501, Biolegend) was coated with 1.2 μg/mL HERV-K113 Capsid protein (sp|Q902F9|90-632, Genescript) diluted in ELISA coating buffer (421701, Biolegend) at 4 ℃ overnight. After three-time washes with PBS, the plate was blocked by blocking solution (1% BSA) in PBS for 1 hour at room temperature. Then we washed the plate with PBS three times again. Then, patient samples were diluted (1:100) and incubated in the coated plates for 3 hours at RT. Then HRP-conjugated secondary antibody was added (SSA001, Sino Biological), followed by a 30-minute incubation. At last, TMB reagent (421101, Biolegend) was added to the wells and waited 10 minutes stopped by TMB stop solution (423001, Biolegend). Each well was analyzed for the optical density (OD) with a microplate reader set to 450 nm.

### FACS analysis

Tissue was digested in culture medium supplemented with 0.6 mg/ml collagenase IV (17104019, Gibco) and 0.01 mg/ml DNase I (11284932001, Merck) at 37°C for 1 hour. Digested tissues were passed through a 0.4-µM cell strainer (352340, BD) to obtain single-cell suspensions, followed by centrifugation at 300 g for 5 min and resuspension in 36% Percoll (P4937, Sigma). After another round of centrifugation at 500 g for 5 min, cells were resuspended in 10 ml of blood lysis buffer (555899, BD) for 10 min at RT to remove red blood cells. After red blood cell lysis, cell suspensions were centrifuged again at 300 g for 5 min and then preincubated with 2.5 µg/ml Fc blocker (156604 for mouse, 422302 for human, BioLegend) on ice for 10 min. After that, the suspensions were further incubated with fluorochrome-labeled antibodies (Table. S2) at 4°C for 20 min. The antibody panel is listed in Supplementary Table S2. For analysis, the samples were washed and resuspended in PBS supplemented with 2% FBS for analysis on a five-laser flow cytometer (BD Biosciences, Fortessa). The data were analysed with FlowJo software (TreeStar).

### ROS detection

We used the CellROX™ kit (C10492, Thermofisher) to detect ROS. In terms of the FACS method, liver tissues were digested to singlets firstly as the method described above. Then we incubated dissociated cells with a ROS probe followed by analysis on a flow cytometer. For the tissue imaging to detect ROS, as the method of immunofluorescence, ROS probes were incubated on tissue slides. After DAPI staining, slides were scanned by confocal microscopy.

### Library preparation

Library preparation, sequencing, and pre-processing service were provided by OE Biotech Co., Ltd (Shanghai, China). Briefly, the countess (Thermo) was used to count the concentration of the single-cell suspension. And the concentration of the single-cell suspension was adjusted to 1000 cells/μL. Cells were loaded to the BD sequencing chip according to the manufacturer’s protocol to capture 5000-10,000 cells/chip position. All the remaining procedures including the library construction were performed according to the standard manufacturer’s protocol.

### scRNA-seq data analysis

Single-cell libraries were sequenced on an Illumina NovaSeq instrument using 150 nt paired-end sequencing. The Cell Ranger 2.1.0 pipeline with the default and recommended parameters were used to process reads. FASTQs generated from Illumina sequencing output were aligned to the human reference genome (hg19) using the STAR algorithm. Gene-Barcode matrices were generated for individual samples by counting unique molecular identifiers (UMIs) and filtering non-cell associated barcodes. Finally, a gene-barcode matrix containing the barcoded cells and gene expression counts were generated. The raw gene expression matrix was loaded into the Seurat package ^20–22^ for quality control and downstream analysis. The cells that contained less than 200 and more than 6000 unique features and contained more than 20% mitochondrial genes were considered as low-quality cells, which were removed for downstream analysis. The high-quality cells were further normalized using the *NormalizedData* function. Top 2000 highly variable genes were found via the *vst* method by *FindVariableFeatures* function. To eliminate the batch effect, gene expression matrices from different samples were integrated based on anchors using function *FindIntegrationAnchors* and *IntegrateData* functions, and the integrated data were further scaled using *ScaleData.*

Principal component analysis (PCA) was performed using scaled integrated data, the top 20 PCs were selected as the input for dimension reduction and cluster. Uniform Manifold Approximation and Projection (UMAP) and graph-based clusters were applied using *RunUMAP, FindNeighbors,* and *FindClusters* function. Top 20 cluster-specific marker genes derived from the *FindAllMarkers* function were employed for manual annotation.

### CyTOF data analysis

CyTOF data was pre-processed with FlowJo to manually gate CD45+ cell population from Flow cytometry standard (FCS) files. The resulting populations were fed to Cytosplore software ^28, 29^ for major cluster identification. Spanning-tree Progression Analysis of Density-normalized Events (SPADE) algorithm was applied on hyperbolic arcsinh the transformed data with a cofactor of 5, for clustering. Hierarchical Stochastic Neighbor Embedding (HSNE) was performed for dimensionality reduction.

The major cluster re-clustering was performed using Unsupervised clustering was performed using the cytofkit package ^70^ in R software. PhenoGraph along with the default parameter was applied for sub-cluster identification. Uniform Manifold Approximation and Projection (UMAP) was employed for dimension reduction. Statistical analysis was performed by the Wilcoxon test, and a p-value less than 0.05 was considered as statistical significance. Visualization was implemented in ggplot2 ^71^.

### scITD analysis

The expression matrix, metadata and selected cell types which at least 3 cells for each sample were processed to form a tensor with dimensions of samples * genes * cell-types. The pseudo-bulked data was normalized following the guideline. The ICA decomposition was performed to analyse the interesting biological variation with recommended parameters. The parameter of top factors and gene sets were determined by stability analysis. The association between samples and group was shown on the top of sample score matrix plot. The corresponding loading matrices for significant factor were generated to determine which genes from each cell type are significant associated with the target factor.

### Lineage tracing analysis

The Bam file for each requested cell was extracted based on the corresponding barcode sequences. Freebayes was applied to call mutations for individual cells and the called mutations were further filtered with minimal depth of two for both reference and allele reads. The lineage profile for each sample was inferred from mutations showing in no more than 80% of all cells and no less than 20% for either Inflammatory or Resident Macrophages. Annovar was subsequently used to annotate filtered mutations.

## Statistical analysis

Raw data obtained from FACS was copied into GraphPad software. Statistical tests were selected based on the appropriate assumptions for the data distribution and variability characteristics. Sample data were analyzed by a two-tailed Student’s t-test to identify statistically significant differences between two groups. One-way ANOVA with the Bonferroni post-test was used to identify differences among three or more groups. The data are represented as the means ± SEMs. A *p*-value < 0.05 indicated statistical significance. The R package “survminer” was implemented to analyze survival differences between different groups.

## Author contributions

Jianpeng Sheng, Junlei Zhang and Yaxing Zhao designed and performed experiments, performed data analysis, and wrote the manuscript, these authors mentioned above contributed equally to this work; Jinyuan Song, Jianghui Tang, Xun Wang, Yongtao Ji, Jiangchao Wu, Taohong Li, and Hui Zhang performed experiments; Sarah Raye Langley and Vincent Tano reviewed the scRNA-seq analysis pipeline and modified the manuscript. Tingbo Liang and Xueli Bai generated the idea, supervised the project, reviewed the manuscript, and secured funding.

## Data Availability

Both the sequencing data and Cy-TOF data were deposited to the Genome Sequence Archive of Beijing Institute of Genomics, Chinese Academy of Sciences with an project accession of PRJCA007722. All data related to this work are set to be accessed upon publication.

## Abbreviation

BA: Biliary atresia
scRNA-seq: Single-cell RNA sequencing
CyTOF: cytometry by time of flight
IHC: immunohistochemistry
mIHC: multiplex IHC
HERV: Human endogenous retrovirus
HP: Human parvovirus
ALT: alanine aminotransferase
AST: aspartate aminotransferase
AchE: acetylcholin esterase
TBIL: total bilirubin
ALP: alkaline phosphatase
γ-GT: γ-glutamyl transpeptidase
UMAP: Uniform Manifold Approximation and Projection
HSNE: Hierarchical Stochastic Neighbor Embedding
ICA: independent components analysis
Teff: T effector cells
Tem: effector memory T cells
Trm: resident T cells
Tcm: central memory T cells
Treg: regulatory T cells
NAC: N acetylcysteine
DTR: Diphtheria toxin receptor
DT: Diphtheria toxin.

## Acknowledgments

The authors would like to thank Lei Ni for mouse management; Qi Chen and Meisheng Zhou for administrative help. This work was supported by the National Key Research and Development Program of China (grant 2019YFA0803000 to J.S.), the Excellent Youth Foundation of Zhejiang Scientific (grant R22H1610037 to J.S.), the National Natural Science Foundation of China (grant 82173078 to J.S.), the Natural Science Foundation of Zhejiang Province (grant 2022C03037 to J.S.), the National Natural Science Foundation of China (grant 81871925 to X.B and grant 82188102 to T.L.), the National Key Research and Development Program (grant 2019YFC1316000 to T.L.) and the National Natural Science Foundation Basic Science Centre of China (Study of Tumour Material and Energy Dynamics, 8218810).

---

**Figure S1.**
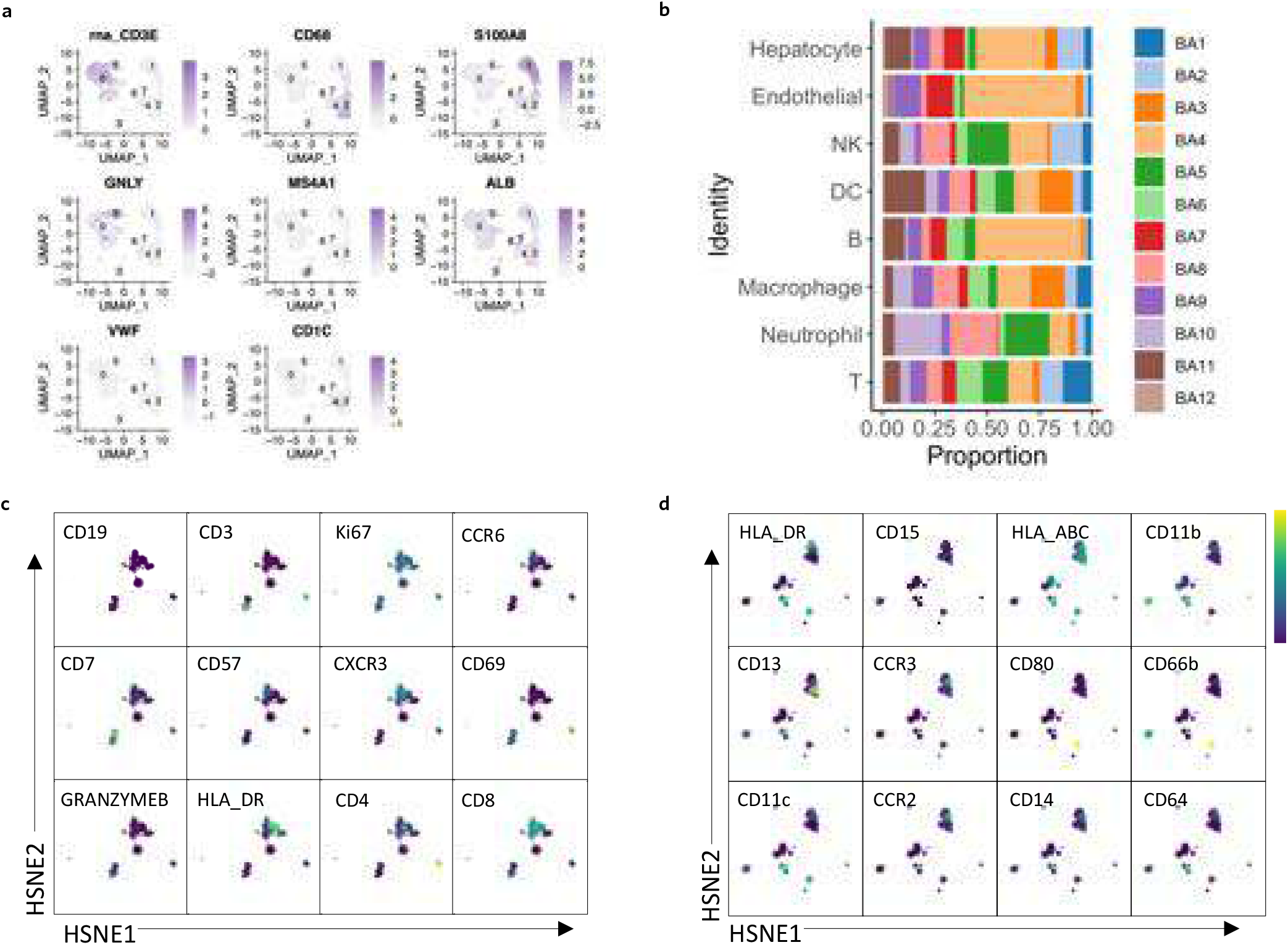
The expression of major cell type markers for scRNA-seq and CyTOF data. **A.** UMAP plots of scRNA-seq data highlighting the expression of marker genes for the main cell types, including T cells (CD3E), macrophage (CD68), neutrophiles (S100A8), NK cells (GNLY), B cells (MS4A1), hepatocytes (ALB), endothelial cells (VWF) and DC (CD1c). **B.** Distribution of major cell types defined by scRNA-seq among different patients. **C.** HSNE plots of the lymphoid panel of CyTOF data highlighting the expression of marker genes for the main cell types, including T cells (CD3, CD4, CD8), B cells (CD19), and NK cells (CD57, GRANZYME B). **D.** HSNE plots of the myeloid panel of CyTOF data highlighting the expression of marker genes for the main cell types, including macrophage (CD68, CD206, and CD86), granulocyte (CD15 and CD66b), and DC (CD1c).

**Figure S2.**
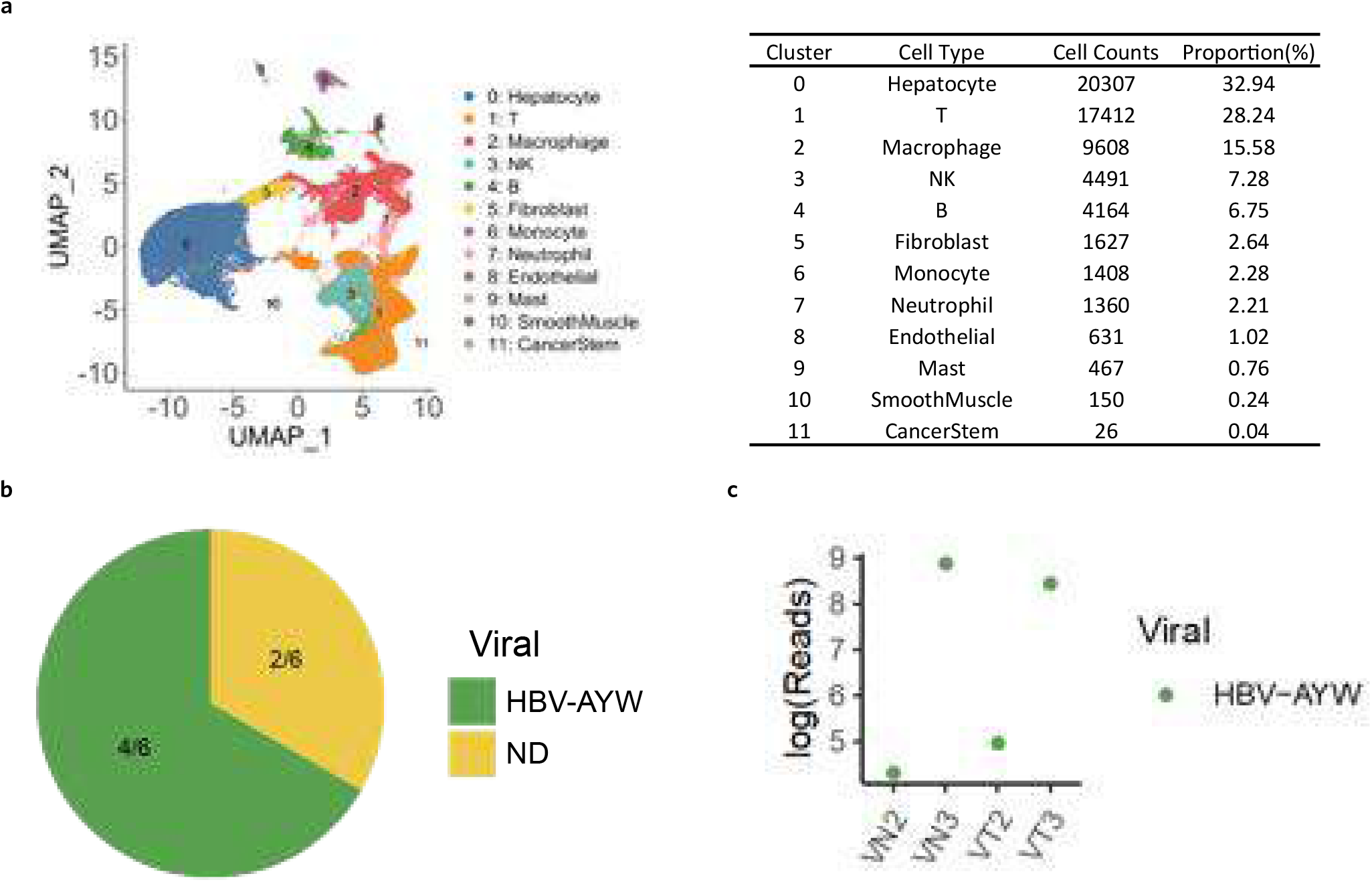
Virus detection in HCC patients. **A.** 12 major cell populations were defined by scRNA-seq analysis based on 6 HCC samples (3 pairs of tumour, VT, and their para-tumour control, VN), shown by UMAP. And cell count for each population was also calculated after quality control. **B.** Pie chart showing the infection map of HCC patients. HBV-AYW was detected in 4 out of 6 HCC patient samples (2 tumour sites and 2 corresponding para-tumour site, green). Viral-Track didn’t spot any virus infection in the remaining 2 patients (yellow). **C.** Viral reads in infected patients. Viral reads shown in log form were calculated for all the virus-infected HCC patient samples. The green colour indicated the HBV-AYW virus.

**Figure S3.**
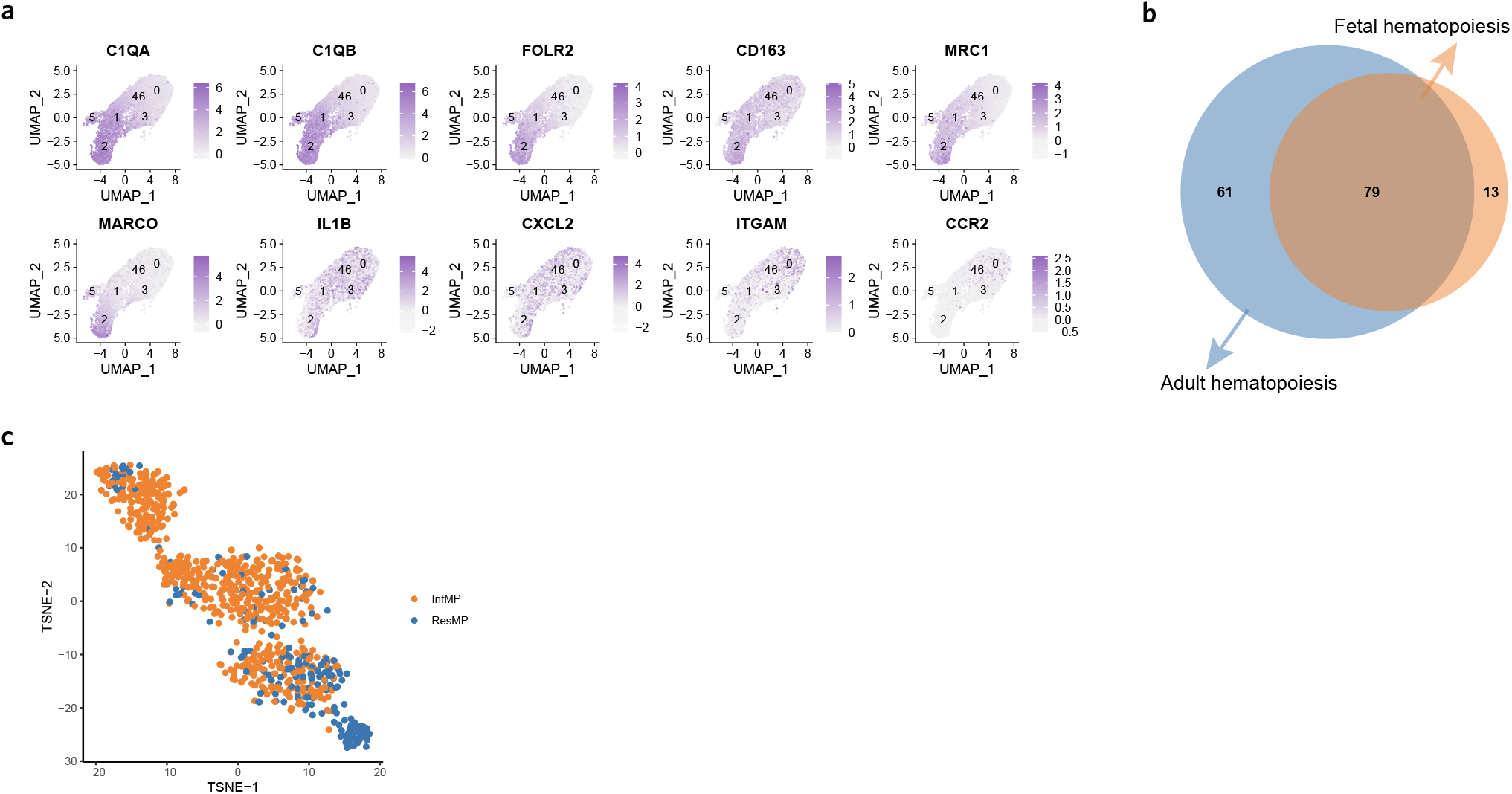
Genetic lineage tracing of macrophages. **A.** UMAP plots of macrophage population highlighting the expression of marker genes for the refined cell clusters, including C1QA, C1QB, FOLR2, CD163, MRC1, MARCO, IL1B, CXCL2, CXCL8, CD11b (encoded by ITGAM), and CCR2. **B and C.** Venn and tSNE plots of mutational profiles for macrophages. Mitochondria and genomic mutations were extracted for each macrophage for each patient and one of the patient macrophage mutational profiles was shown.

**Figure S4.**
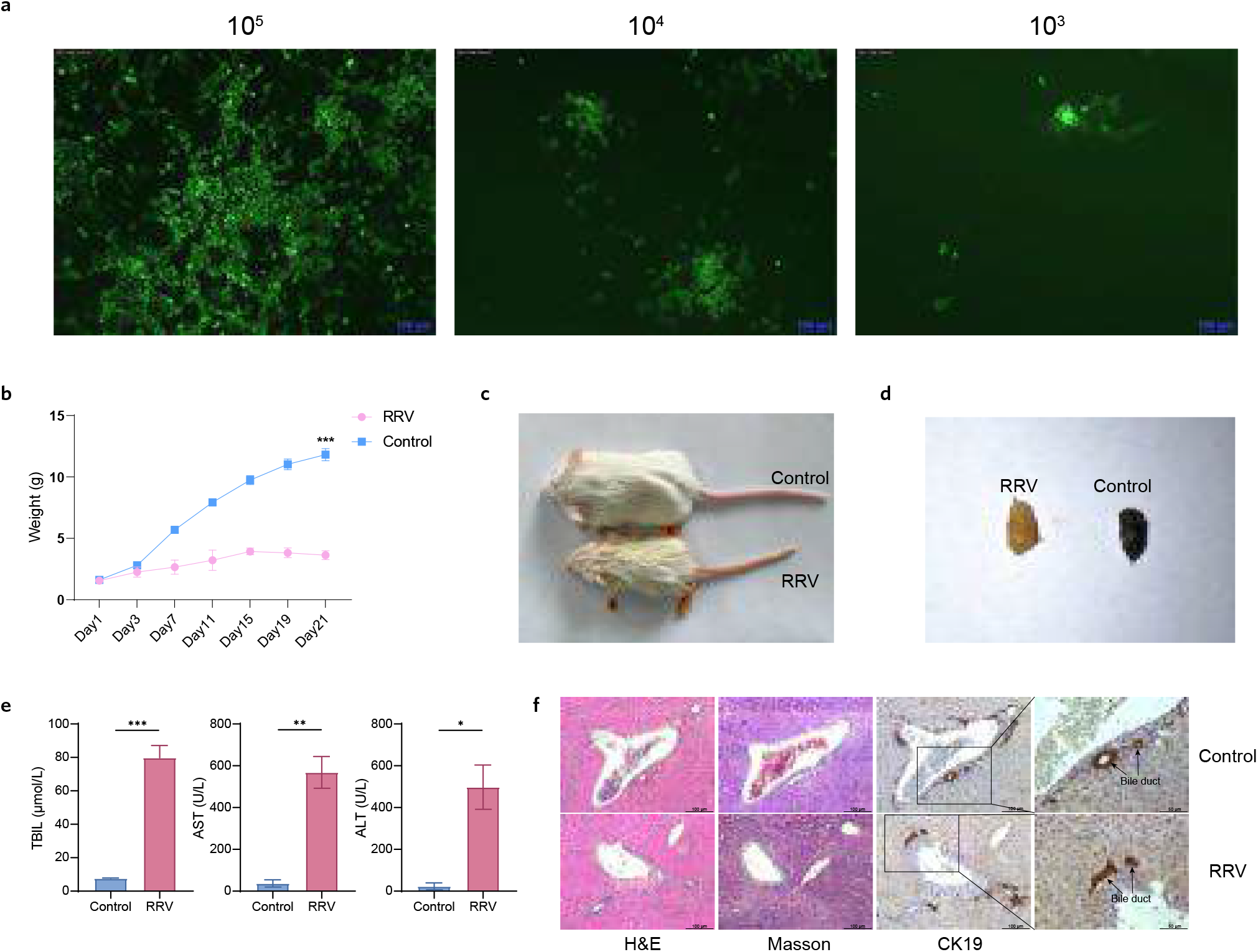
The establishment of the RRV-infected mouse model. **A.** Immunofluorescent staining images show that RRV with different concentrations of 10^5^, 10^4,^ and 10^3^ FFU/mL infected the MA104 cell line. **B.** The weight change of mice infected with RRV compared to group control. **C.** The size, skin, and mucous membrane change between the two groups. RRV mouse is smaller and its skin and mucous become yellow. **D.** Feces of RRV mice become yellow compared to group control. **E.** Total bilirubin, AST, and ALT in serum from group RRV increase dramatically. The images of H&E, Masson, and IHC staining show the pathological change between the two groups.

| Number | Age | Gender | Diagnosis | Date of surgery | ALT (U/L) | AST (U/L) |
| --- | --- | --- | --- | --- | --- | --- |
| 1 | 4M8D | F | Biliary Atresia | 2021/3/15 | 196 | 238 |
| 2 | 4M20D | F | Biliary Atresia | 2021/3/25 | 173 | 284 |
| 3 | 5M8D | M | Biliary Atresia | 2021/3/27 | 260 | 241 |
| 4 | 4M12D | M | Biliary Atresia | 2021/5/11 | 121 | 373 |
| 5 | 5M4D | F | Biliary Atresia | 2021/5/11 | 140 | 201 |
| 6 | 5M5D | M | Biliary Atresia | 2021/5/14 | 194 | 442 |
| 7 | 5M9D | M | Biliary Atresia | 2021/5/25 | 179 | 168 |
| 8 | 9M11D | F | Biliary Atresia | 2021/6/22 | 68 | 252 |
| 9 | 7M4D | F | Biliary Atresia | 2021/6/30 | 146 | 351 |
| 10 | 7M15D | M | Biliary Atresia | 2021/6/29 | 391 | 474 |
| 11 | 7M7D | F | Biliary Atresia | 2021/6/29 | 107 | 232 |
| 12 | 7M14D | F | Biliary Atresia | 2021/6/16 | 238 | 277 |
| Mean | 6.1M |  |  |  | 184 | 294 |
| SD | 1.59M |  |  |  | 85 | 95 |

The information of patients (ELISA)
| Number | Age | Gender | Diagnosis | Date of surgery | With father | With mother |
| --- | --- | --- | --- | --- | --- | --- |
| 1 | 5M18D | F | Biliary Atresia | 2021/8/9 | √ | √ |
| 2 | 5M6D | M | Biliary Atresia | 2021/8/9 | √ | √ |
| 3 | 4M28D | F | Biliary Atresia | 2021/8/11 | √ | √ |
| 4 | 6M1D | M | Biliary Atresia | 2021/8/17 | √ | √ |
| 5 | 5M27D | F | Biliary Atresia | 2021/8/20 | √ | √ |
| 6 | 4M29D | F | Biliary Atresia | 2021/8/25 | √ | √ |
| 7 | 6M8D | M | Biliary Atresia | 2021/8/31 | √ | √ |
| 8 | 9M4D | M | Biliary Atresia | 2021/8/31 | √ | √ |
| 9 | 7M27D | M | Biliary Atresia | 2021/9/6 | √ | √ |
| 10 | 7M | F | Biliary Atresia | 2021/9/14 | √ | √ |
| 11 | 3M4D | F | Biliary Atresia | 2021/9/14 | √ | × |
| 12 | 5M23D | F | Biliary Atresia | 2021/9/15 | √ | √ |
| 13 | 9M | M | Biliary Atresia | 2021/9/20 | √ | √ |
| 14 | 8M3D | F | Biliary Atresia | 2021/9/28 | √ | √ |
| 15 | 3M5D | M | Biliary Atresia | 2021/9/29 | √ | √ |
| 16 | 5M17D | F | Biliary Atresia | 2021/10/6 | √ | √ |

f)
| AchE (U/L) | TBIL (μmol/L) | ALP (U/L) | γ-GT (U/L) |
| --- | --- | --- | --- |
| 5179 | 132.5 | 532 | 447 |
| 3712 | 218.2 | 754 | 617 |
| 2229 | 78.2 | 265 | 33 |
| 1459 | 367.7 | 535 | 50 |
| 2145 | 192.4 | 403 | 488 |
| 1503 | 501.8 | 987 | 57 |
| 6292 | 206.5 | 547 | 894 |
| 3265 | 139.2 | 281 | 389 |
| 1651 | 209.6 | 272 | 260 |
| 3161 | 189.3 | 277 | 1573 |
| 2830 | 270.4 | 314 | 198 |
| 2755 | 238.9 | 271 | 572 |
| 3015 | 229 | 453 | 465 |
| 1478 | 113 | 230 | 436 |

FACS Antibody Panel
| Species | Antibodies | clone | source | Cat# |
| --- | --- | --- | --- | --- |
| human | CD45 | HI30 | BD | 563716 |
| human | CD16 | B7311 | BD | 561313 |
| human | CD11b | 1CRF44 | BD | 562721 |
| human | CD68 | Y1/82A | Biolegend | 333808 |
| human | CD56 | 5.1H11 | Biolegend | 362510 |
| human | Granzyme B | QA18A28 | Biolegend | 396414 |
| human | CD57 | HNK-1 | Biolegend | 359604 |
| human | CD158a | HP-DM1 | Biolegend | 374903 |
| human | CD158d | mAb 33 (33) | Biolegend | 347007 |
| mouse | CD45 | 30-F11 | BD | 563053 |
| mouse | CD11b | M1/70 | BD | 562287 |
| mouse | MHCII (I-A/I-E) | M5/114.15.2 | BD | 562564 |
| mouse | F4/80 | BM8 | Biolegend | 123114 |
| mouse | Ly-6C | HK1.4 | Biolegend | 128036 |
| mouse | Ly-6G | 1A8 | Biolegend | 127605 |

## Reference

1 Hartley, J. L., Davenport, M. & Kelly, D. A. Biliary atresia. Lancet 374, 1704–1713, doi:10.1016/S0140-6736(09)60946-6 (2009).

2 Murase, N. et al. A New Era of Laparoscopic Revision of Kasai Portoenterostomy for the Treatment of Biliary Atresia. Biomed Res Int 2015, 173014, doi:10.1155/2015/173014 (2015).

3 Landing, B. H. Considerations of the pathogenesis of neonatal hepatitis, biliary atresia and choledochal cyst--the concept of infantile obstructive cholangiopathy. Prog Pediatr Surg 6, 113–139 (1974).

4 Chang, M. H. et al. Polymerase chain reaction to detect human cytomegalovirus in livers of infants with neonatal hepatitis. Gastroenterology 103, 1022–1025, doi:10.1016/0016-5085(92)90038-z (1992).

5 Xu, Y. et al. The perinatal infection of cytomegalovirus is an important etiology for biliary atresia in China. Clin Pediatr (Phila) 51, 109–113, doi:10.1177/0009922811406264 (2012).

6 Zani, A., Quaglia, A., Hadzic, N., Zuckerman, M. & Davenport, M. Cytomegalovirus-associated biliary atresia: An aetiological and prognostic subgroup. J Pediatr Surg 50, 1739–1745, doi:10.1016/j.jpedsurg.2015.03.001 (2015).

7 Weaver, L. T., Nelson, R. & Bell, T. M. The association of extrahepatic bile duct atresia and neonatal Epstein-Barr virus infection. Acta Paediatr Scand 73, 155–157, doi:10.1111/j.1651-2227.1984.tb09918.x (1984).

8 Kikuchi, K. et al. Vanishing bile duct syndrome associated with chronic EBV infection. Dig Dis Sci 45, 160–165, doi:10.1023/a:1005434015863 (2000).

9 Fjaer, R. B., Bruu, A. L. & Nordbo, S. A. Extrahepatic bile duct atresia and viral involvement. Pediatr Transplant 9, 68–73, doi:10.1111/j.1399-3046.2005.00257.x (2005).

10 Bobo, L. et al. Lack of evidence for rotavirus by polymerase chain reaction/enzyme immunoassay of hepatobiliary samples from children with biliary atresia. Pediatr Res 41, 229–234, doi:10.1203/00006450-199702000-00013 (1997).

11 Esona, M. D., Humphrey, C. D., Dennehy, P. H. & Jiang, B. Prevalence of group C rotavirus among children in Rhode Island, United States. J Clin Virol 42, 221–224, doi:10.1016/j.jcv.2008.02.002 (2008).

12 Rauschenfels, S. et al. Incidence of hepatotropic viruses in biliary atresia. Eur J Pediatr 168, 469–476, doi:10.1007/s00431-008-0774-2 (2009).

13 Saito, T. et al. Lack of evidence for reovirus infection in tissues from patients with biliary atresia and congenital dilatation of the bile duct. J Hepatol 40, 203–211, doi:10.1016/j.jhep.2003.10.025 (2004).

14 Saito, T. et al. Evidence for viral infection as a causative factor of human biliary atresia. J Pediatr Surg 50, 1398–1404, doi:10.1016/j.jpedsurg.2015.04.006 (2015).

15 Averbukh, L. D. & Wu, G. Y. Evidence for Viral Induction of Biliary Atresia: A Review. J Clin Transl Hepatol 6, 410–419, doi:10.14218/JCTH.2018.00046 (2018).

16 Kolodziejczyk, A. A., Kim, J. K., Svensson, V., Marioni, J. C. & Teichmann, S. A. The technology and biology of single-cell RNA sequencing. Mol Cell 58, 610–620, doi:10.1016/j.molcel.2015.04.005 (2015).

17 Bandura, D. R. et al. Mass cytometry: technique for real time single cell multitarget immunoassay based on inductively coupled plasma time-of-flight mass spectrometry. Anal Chem 81, 6813–6822, doi:10.1021/ac901049w (2009).

18 Wang, J. et al. Liver Immune Profiling Reveals Pathogenesis and Therapeutics for Biliary Atresia. Cell 183, 1867–1883 e1826, doi:10.1016/j.cell.2020.10.048 (2020).

19 Bost, P. et al. Host-Viral Infection Maps Reveal Signatures of Severe COVID-19 Patients. Cell 181, 1475–1488 e1412, doi:10.1016/j.cell.2020.05.006 (2020).

20 Hao, Y. et al. Integrated analysis of multimodal single-cell data. Cell 184, 3573–3587 e3529, doi:10.1016/j.cell.2021.04.048 (2021).

21 Stuart, T. et al. Comprehensive Integration of Single-Cell Data. Cell 177, 1888–1902 e1821, doi:10.1016/j.cell.2019.05.031 (2019).

22 Butler, A., Hoffman, P., Smibert, P., Papalexi, E. & Satija, R. Integrating single-cell transcriptomic data across different conditions, technologies, and species. Nat Biotechnol 36, 411–420, doi:10.1038/nbt.4096 (2018).

23 Becht, E. et al. Dimensionality reduction for visualizing single-cell data using UMAP. Nat Biotechnol, doi:10.1038/nbt.4314 (2018).

24 Chinzei, R. et al. Embryoid-body cells derived from a mouse embryonic stem cell line show differentiation into functional hepatocytes. Hepatology 36, 22–29, doi:10.1053/jhep.2002.34136 (2002).

25 Vischer, U. M. von Willebrand factor, endothelial dysfunction, and cardiovascular disease. J Thromb Haemost 4, 1186–1193, doi:10.1111/j.1538-7836.2006.01949.x (2006).

26 van Unen, V. et al. Visual analysis of mass cytometry data by hierarchical stochastic neighbour embedding reveals rare cell types. Nat Commun 8, 1740, doi:10.1038/s41467-017-01689-9 (2017).

27 Boller, K. et al. Human endogenous retrovirus HERV-K113 is capable of producing intact viral particles. J Gen Virol 89, 567–572, doi:10.1099/vir.0.83534-0 (2008).

28 Sokal, E. M. et al. Acute parvovirus B19 infection associated with fulminant hepatitis of favourable prognosis in young children. Lancet 352, 1739–1741, doi:10.1016/S0140-6736(98)06165-0 (1998).

29 Jonathan Mitchel, E. B., and Peter Kharchenko. scITD: Single-Cell Interpretable Tensor Decomposition. R package version 1.*0*.*1* (2021).

30 Duurland, C. L. et al. CD161 expression and regulation defines rapidly responding effector CD4+ T cells associated with improved survival in HPV16-associated tumors. J Immunother Cancer 10, doi:10.1136/jitc-2021-003995 (2022).

31 Martin, M. D. & Badovinac, V. P. Defining Memory CD8 T Cell. Front Immunol 9, 2692, doi:10.3389/fimmu.2018.02692 (2018).

32 Herndler-Brandstetter, D. et al. KLRG1(+) Effector CD8(+) T Cells Lose KLRG1, Differentiate into All Memory T Cell Lineages, and Convey Enhanced Protective Immunity. Immunity 48, 716–729 e718, doi:10.1016/j.immuni.2018.03.015 (2018).

33 Eberl, M. et al. Human Vgamma9/Vdelta2 effector memory T cells express the killer cell lectin-like receptor G1 (KLRG1). J Leukoc Biol 77, 67–70, doi:10.1189/jlb.0204096 (2005).

34 Schoggins, J. W. & Randall, G. Lipids in innate antiviral defense. Cell Host Microbe 14, 379–385, doi:10.1016/j.chom.2013.09.010 (2013).

35 Zhao, J., Chen, J., Li, M., Chen, M. & Sun, C. Multifaceted Functions of CH25H and 25HC to Modulate the Lipid Metabolism, Immune Responses, and Broadly Antiviral Activities. Viruses 12, doi:10.3390/v12070727 (2020).

36 Yeh, J. H., Sidhu, S. S. & Chan, A. C. Regulation of a late phase of T cell polarity and effector functions by Crtam. Cell 132, 846–859, doi:10.1016/j.cell.2008.01.013 (2008).

37 Xiong, Q. et al. Characteristics of SARS-CoV-2-specific cytotoxic T cells revealed by single-cell immune profiling of longitudinal COVID-19 blood samples. Signal Transduct Target Ther 5, 285, doi:10.1038/s41392-020-00425-y (2020).

38 Zaiss, D. M. et al. Amphiregulin enhances regulatory T cell-suppressive function via the epidermal growth factor receptor. Immunity 38, 275–284, doi:10.1016/j.immuni.2012.09.023 (2013).

39 Manes, T. D. & Pober, J. S. Identification of endothelial cell junctional proteins and lymphocyte receptors involved in transendothelial migration of human effector memory CD4+ T cells. J Immunol 186, 1763–1768, doi:10.4049/jimmunol.1002835 (2011).

40 Best, J. A. et al. Transcriptional insights into the CD8(+) T cell response to infection and memory T cell formation. Nat Immunol 14, 404–412, doi:10.1038/ni.2536 (2013).

41 Hamann, I. et al. Analyses of phenotypic and functional characteristics of CX3CR1-expressing natural killer cells. Immunology 133, 62–73, doi:10.1111/j.1365-2567.2011.03409.x (2011).

42 Ponti, C. et al. Role of CREB transcription factor in c-fos activation in natural killer cells. Eur J Immunol 32, 3358–3365, doi:10.1002/1521-4141(200212)32:12<3358::AID-IMMU3358>3.0.CO;2-Q (2002).

43 Russick, J. et al. Natural killer cells in the human lung tumor microenvironment display immune inhibitory functions. J Immunother Cancer 8, doi:10.1136/jitc-2020-001054 (2020).

44 He, H. et al. Single-cell transcriptome analysis of human skin identifies novel fibroblast subpopulation and enrichment of immune subsets in atopic dermatitis. J Allergy Clin Immunol 145, 1615–1628, doi:10.1016/j.jaci.2020.01.042 (2020).

45 Cobaleda, C., Schebesta, A., Delogu, A. & Busslinger, M. Pax5: the guardian of B cell identity and function. Nat Immunol 8, 463–470, doi:10.1038/ni1454 (2007).

46 Maghazachi, A. A. Role of chemokines in the biology of natural killer cells. Curr Top Microbiol Immunol 341, 37–58, doi:10.1007/82_2010_20 (2010).

47 Fensterl, V. et al. Interferon-induced Ifit2/ISG54 protects mice from lethal VSV neuropathogenesis. PLoS Pathog 8, e1002712, doi:10.1371/journal.ppat.1002712 (2012).

48 Spaan, M. et al. Immunological Analysis During Interferon-Free Therapy for Chronic Hepatitis C Virus Infection Reveals Modulation of the Natural Killer Cell Compartment. J Infect Dis 213, 216–223, doi:10.1093/infdis/jiv391 (2016).

49 Cho, H., Shrestha, B., Sen, G. C. & Diamond, M. S. A role for Ifit2 in restricting West Nile virus infection in the brain. J Virol 87, 8363–8371, doi:10.1128/JVI.01097-13 (2013).

50 Wiedemann, G. M., Geary, C. D., Lau, C. M. & Sun, J. C. Cutting Edge: STAT1-Mediated Epigenetic Control of Rsad2 Promotes Clonal Expansion of Antiviral NK Cells. J Immunol 205, 21–25, doi:10.4049/jimmunol.2000086 (2020).

51 Rivera-Serrano, E. E. et al. Viperin Reveals Its True Function. Annu Rev Virol 7, 421–446, doi:10.1146/annurev-virology-011720-095930 (2020).

52 Ali, A. et al. Natural killer cell immunosuppressive function requires CXCR3-dependent redistribution within lymphoid tissues. J Clin Invest 131, doi:10.1172/JCI146686 (2021).

53 Sordo-Bahamonde, C. et al. BTLA/HVEM Axis Induces NK Cell Immunosuppression and Poor Outcome in Chronic Lymphocytic Leukemia. Cancers (Basel) 13, doi:10.3390/cancers13081766 (2021).

54 Harizi, H. Reciprocal crosstalk between dendritic cells and natural killer cells under the effects of PGE2 in immunity and immunopathology. Cell Mol Immunol 10, 213–221, doi:10.1038/cmi.2013.1 (2013).

55 Gordon, S. M. et al. The transcription factors T-bet and Eomes control key checkpoints of natural killer cell maturation. Immunity 36, 55–67, doi:10.1016/j.immuni.2011.11.016 (2012).

56 van den Bosch, M. H. et al. Alarmin S100A9 Induces Proinflammatory and Catabolic Effects Predominantly in the M1 Macrophages of Human Osteoarthritic Synovium. J Rheumatol 43, 1874–1884, doi:10.3899/jrheum.160270 (2016).

57 Sanjurjo, L. et al. CD5L Promotes M2 Macrophage Polarization through Autophagy-Mediated Upregulation of ID3. Front Immunol 9, 480, doi:10.3389/fimmu.2018.00480 (2018).

58 Mahabeleshwar, G. H. et al. The myeloid transcription factor KLF2 regulates the host response to polymicrobial infection and endotoxic shock. Immunity 34, 715–728, doi:10.1016/j.immuni.2011.04.014 (2011).

59 Tang, J., Shi, Y., Zhan, L. & Qin, C. Downregulation of GPR183 on infection restricts the early infection and intracellular replication of mycobacterium tuberculosis in macrophage. Microb Pathog 145, 104234, doi:10.1016/j.micpath.2020.104234 (2020).

60 Yang, J. et al. Naringenin inhibits proinflammatory cytokine production in macrophages through inducing MT1G to suppress the activation of NFkappaB. Mol Immunol 137, 155–162, doi:10.1016/j.molimm.2021.07.003 (2021).

61 Wu, X. et al. Single-cell sequencing of immune cells from anticitrullinated peptide antibody positive and negative rheumatoid arthritis. Nat Commun 12, 4977, doi:10.1038/s41467-021-25246-7 (2021).

62 Asano, K. et al. Intestinal CD169(+) macrophages initiate mucosal inflammation by secreting CCL8 that recruits inflammatory monocytes. Nat Commun 6, 7802, doi:10.1038/ncomms8802 (2015).

63 Sheng, J., Ruedl, C. & Karjalainen, K. Most Tissue-Resident Macrophages Except Microglia Are Derived from Fetal Hematopoietic Stem Cells. Immunity 43, 382–393, doi:10.1016/j.immuni.2015.07.016 (2015).

64 Sheng, J., Ruedl, C. & Karjalainen, K. Fetal HSCs versus EMP2s. Immunity 43, 1025, doi:10.1016/j.immuni.2015.11.023 (2015).

65 Sheng, J. et al. Topological analysis of hepatocellular carcinoma tumour microenvironment based on imaging mass cytometry reveals cellular neighbourhood regulated reversely by macrophages with different ontogeny. Gut, gutjnl-2021-324339, doi:10.1136/gutjnl-2021-324339 (2021).

66 Sheng, J. et al. Fate mapping analysis reveals a novel murine dermal migratory Langerhans-like cell population. Elife 10, doi:10.7554/eLife.65412 (2021).

67 Ludwig, L. S. et al. Lineage Tracing in Humans Enabled by Mitochondrial Mutations and Single-Cell Genomics. Cell 176, 1325–1339 e1322, doi:10.1016/j.cell.2019.01.022 (2019).

68 Sokulsky, L. A. et al. A Critical Role for the CXCL3/CXCL5/CXCR2 Neutrophilic Chemotactic Axis in the Regulation of Type 2 Responses in a Model of Rhinoviral-Induced Asthma Exacerbation. J Immunol 205, 2468–2478, doi:10.4049/jimmunol.1901350 (2020).

69 Jalal, P. J. et al. Expression of human ficolin-2 in hepatocytes confers resistance to infection by diverse hepatotropic viruses. J Med Microbiol 68, 642–648, doi:10.1099/jmm.0.000935 (2019).

70 Huang, K. et al. Long non-coding RNAs (lncRNAs) NEAT1 and MALAT1 are differentially expressed in severe COVID-19 patients: An integrated single-cell analysis. PLoS One 17, e0261242, doi:10.1371/journal.pone.0261242 (2022).

71 Vemula, S. V. et al. Identification of proximal biomarkers of PKC agonism and evaluation of their role in HIV reactivation. Antiviral Res 139, 161–170, doi:10.1016/j.antiviral.2016.11.014 (2017).

72 McCartney, E. M. et al. Alcohol metabolism increases the replication of hepatitis C virus and attenuates the antiviral action of interferon. J Infect Dis 198, 1766–1775, doi:10.1086/593216 (2008).

73 Fabriek, B. O., Dijkstra, C. D. & van den Berg, T. K. The macrophage scavenger receptor CD163. Immunobiology 210, 153–160, doi:10.1016/j.imbio.2005.05.010 (2005).

74 Sheng, J. et al. A Discrete Subset of Monocyte-Derived Cells among Typical Conventional Type 2 Dendritic Cells Can Efficiently Cross-Present. Cell Rep 21, 1203–1214, doi:10.1016/j.celrep.2017.10.024 (2017).

75 Scapini, P. et al. The neutrophil as a cellular source of chemokines. Immunol Rev 177, 195–203, doi:10.1034/j.1600-065x.2000.17706.x (2000).

76 Pias, E. K. et al. Differential effects of superoxide dismutase isoform expression on hydroperoxide-induced apoptosis in PC-12 cells. J Biol Chem 278, 13294–13301, doi:10.1074/jbc.M208670200 (2003).

77 Kilgore, A. & Mack, C. L. Update on investigations pertaining to the pathogenesis of biliary atresia. Pediatr Surg Int 33, 1233–1241, doi:10.1007/s00383-017-4172-6 (2017).

78 Boudreau, J. E. & Hsu, K. C. Natural Killer Cell Education and the Response to Infection and Cancer Therapy: Stay Tuned. Trends Immunol 39, 222–239, doi:10.1016/j.it.2017.12.001 (2018).

79 Gooneratne, S. L., Center, R. J., Kent, S. J. & Parsons, M. S. Functional advantage of educated KIR2DL1(+) natural killer cells for anti-HIV-1 antibody-dependent activation. Clin Exp Immunol 184, 101–109, doi:10.1111/cei.12752 (2016).

